# Spatially mapped single-cell chromatin accessibility

**DOI:** 10.1101/815720

**Authors:** Casey A. Thornton, Ryan M. Mulqueen, Andrew Nishida, Kristof A. Torkenczy, Eve G. Lowenstein, Andrew J. Fields, Frank J. Steemers, Wenri Zhang, Heather L. McConnell, Randy L. Woltjer, Anusha Mishra, Kevin M. Wright, Andrew C. Adey

## Abstract

High-throughput single-cell epigenomic assays can resolve the heterogeneity of cell types and states in complex tissues, however, spatial orientation within the network of interconnected cells is lost. Here, we present a novel method for highly scalable, spatially resolved, single-cell profiling of chromatin states. We use high-density multiregional sampling to perform single-cell combinatorial indexing on Microbiopsies Assigned to Positions for the Assay for Transposase Accessible Chromatin (sciMAP-ATAC) to produce single-cell data of an equivalent quality to non-spatially resolved single-cell ATAC-seq, where each cell is localized to a three-dimensional position within the tissue. A typical experiment comprises between 96 and 384 spatially mapped tissue positions, each producing 10s to over 100 individual single-cell ATAC-seq profiles, and a typical resolution of 214 cubic microns; with the ability to tune the resolution and cell throughput to suit each target application. We apply sciMAP-ATAC to the adult mouse primary somatosensory cortex, where we profile cortical lamination and demonstrate the ability to analyze data from a single tissue position or compare a single cell type in adjacent positions. We also profile the human primary visual cortex, where we produce spatial trajectories through the cortex. Finally, we characterize the spatially progressive nature of cerebral ischemic infarct in the mouse brain using a model of transient middle cerebral artery occlusion. We leverage the spatial information to identify novel and known transcription factor activities that vary by proximity to the ischemic infarction core with cell type specificity.

## Introduction

Heterogeneous cell types coordinate in complex networks to generate emergent properties of tissues. These cell types are not evenly dispersed across tissues, allowing for spatially localized functionality of organs. In many disease states, this becomes more apparent, as the affected organ experiences spatially progressive etiologies. For example, in the case of cerebral ischemia, loss of blood flow due to a blocked artery or blood vessel causes local cell death or infarction. Following ischemic injury, astrocytes and microglia enter reactive states with novel functions that are metered by proximity to the site of infarction^1^, but this spatial information has, so far, been difficult to assess. With the advent of massively parallel sequencing, many methods have emerged that characterize the molecular profiles of cells in unperturbed and perturbed systems by sequencing transcriptomic, genomic, and epigenomic cellular content. Single-cell technologies have further advanced these efforts by enabling the isolation of signals from individual cells within a sample, thus resolving the heterogeneity of complex tissues. Applications of single-cell technologies have identified novel cell types with characteristic-omic signatures in the highly complex tissue of the brain^2,3^. In the cerebral cortex, specifically, cells form an intricate layered hierarchical structure comprised of both neuronal and glial cell types that generate sensory, motor and associational percepts^4^. While single-cell technologies have revolutionized cellular taxonomy and quantification of tissue heterogeneity, the spatial localization of single cells goes unrecorded because of whole-sample dissociation. Layer-specific gene expression profiles of cortical neurons and astrocytes have been characterized by spatial transcriptomic approaches and immunohistochemical (IHC) staining; however, spatially mapped epigenetic states of cortical cells have yet to be directly assayed, without relying on data integration^5–7^.

To address this challenge, several strategies have emerged to assay transcription either directly *in situ* or in a regional manner. The former techniques utilize fluorescence *in situ* hybridization (FISH)^8–10^ or *in situ* RNA-sequencing^11,12^. While powerful, FISH methods require the use of a defined probe set and are limited to the identification of DNA and RNA sequences. In contrast, technologies that utilize array-based mRNA barcoding do not require a defined set of genes and operate similarly to single-cell RNA-seq methods^13,14^, thus allowing for whole transcriptome profiling. Initial iterations of these platforms capture regional transcription over multiple cells; however, higher resolution variants may facilitate single-cell resolution. Unfortunately, these platforms rely on the relatively easy access to mRNA molecules that can be released from the cytoplasm and hybridized to barcoding probes, making the expansion into nuclear epigenetic properties challenging. It is for this reason that none of the current barcode array technologies have yet expanded into the chromatin accessibility space. With the wealth of epigenetic information that resides in the nucleus and the value it can add to characterizing complex biological systems^15–17^, we sought to address this challenge by harnessing the inherent throughput characteristics of single-cell combinatorial indexing assays^18,19^.

Here, we present single-cell combinatorial indexing from Microbiopsies with Assigned Positions for the Assay for Transposase Accessible Chromatin (sciMAP-ATAC). sciMAP-ATAC preserves cellular localization within intact tissues and generates thousands of spatially resolved high quality single-cell ATAC-seq libraries. As with other ‘sci-’ technologies, sciMAP-ATAC does not require specialized equipment and scales nonlinearly, enabling high throughput potential. Building upon multiregional sampling strategies^20,21^, where several regions are isolated, we reasoned that the sample multiplexing capabilities of combinatorial indexing could be utilized to perform high throughput multiregional sampling at resolutions approaching those of array-based spatial transcriptional profiling, all while retaining true single-cell profiles. Unlike multiregional sampling, we perform high-density microbiopsy sampling, ranging from 100-500 µm in diameter, on cryosectioned tissue sections, between 100-300 µm in thickness, to produce up to hundreds of spatially mapped punches of tissue, each producing a set of single-cell chromatin accessibility profiles (Figure 1a). We demonstrate the utility of sciMAP-ATAC by profiling the murine and human cortex, where distinct cell type compositions and chromatin profiles are observed based on the spatial orientation of the punches, and further extend the platform to characterize cerebral ischemic injury in a mouse model system where cell type compositions and epigenetic states are metered by proximity to the injury site (Extended Data Figure 1).

**Figure 1.**
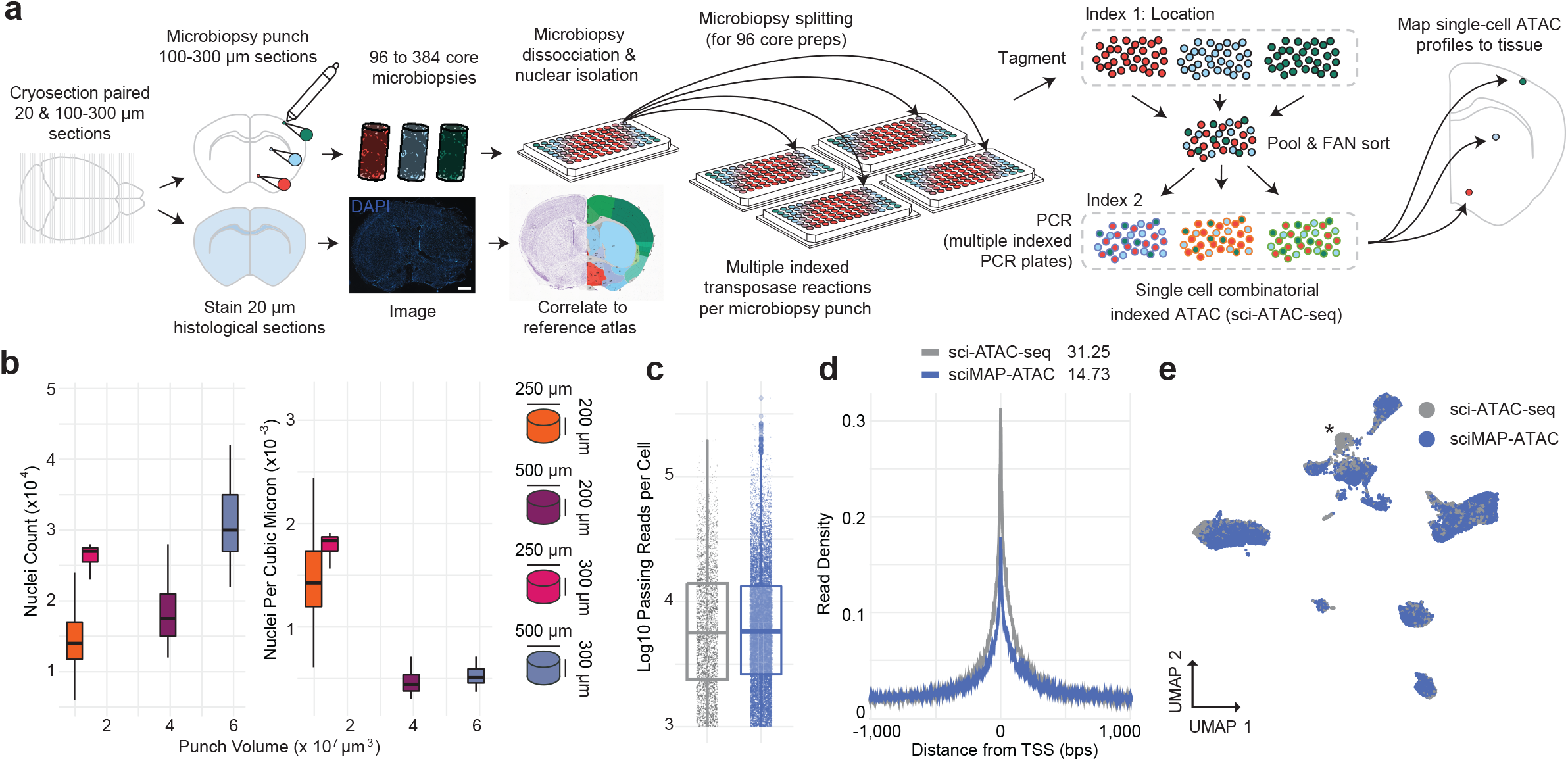
sciMAP-ATAC schematic and performance. **a.** sciMAP-ATAC workflow. Cryosectioning of alternating 20 μm (histological) and 100-300 μm (sciMAP-ATAC) slices are obtained. Thin (20 μm) slices are stained and imaged for use in spatial registration (Scale bar, 1mm). Thick (100-300 μm) slices are carried through high-density microbiopsy punching (100-500 μm diameter) in the cryostat chamber. Punches are placed directly into wells of a microwell plate for nuclei isolation and washed prior to splitting into multiple wells for indexed transposition and the sci-ATAC-seq workflow. **b.** Four punch volumes were assessed for nuclei yield using either a 250 μm or 500 μm diameter punch on a 200 μm or 300 μm thick section. Total nuclei isolated for each punch is shown on the left, and normalized for tissue voxel volume on the right, representing the efficiency of extraction from each punch. **c.** Passing reads per cell from sci-ATAC-seq and sciMAP-ATAC, which are comparable at the level of depth sequenced. **d.** ATAC read signal at TSSs for sci-ATAC-seq and sciMAP-ATAC. Enrichment for sci-ATAC-seq is slightly greater than that of sciMAP-ATAC, likely due to increased processing time of isolated nuclei prior to transposition. **e.** sciMAP-ATAC and sci-ATAC-seq libraries from mouse brain group closely together. Asterisk indicates a population of 734 cells, derived from spinal cord, which was not sampled during microbiopsy punching.

## Results

### Single-cell combinatorial indexed ATAC-seq from microbiopsy punches

Single-cell ATAC-seq requires the isolation and processing of nuclei such that the nuclear scaffold remains intact to facilitate library preparation via transposition *in situ*; it also requires that the chromatin structure is maintained to produce a chromatin accessibility signal. We and others have explored methods for tissue preservation that are compatible with single-cell ATAC-seq^18,22^, however, we sought to confirm that these strategies are compatible with freezing techniques used for cryosectioning and immunohistochemical (IHC) staining of tissue. We tested our workflow on mouse whole brain samples by processing one hemisphere using flash-freezing methods designed for tissue freezing medium (TFM) embedding and cryosectioning (Methods) and processing the paired hemisphere as fresh tissue. Our previously established non-spatially resolved sci-ATAC-seq workflow^22^ was performed on both hemispheres, including pooling post-transposition for sorting, PCR amplification, and sequencing. Flash-frozen and fresh nuclei produced nearly identical passing reads per cell at the depth they were sequenced, along with comparable fractions of reads present in a set of aggregate mouse ATAC-seq peaks (FRiS; 0.93 and 0.91 for fresh and frozen, respectively; Extended Data Figure 2a and 2b).

We then explored techniques for cryosectioning flash-frozen TFM-embedded tissue at thicknesses compatible with microbiopsy punching. Typically, cryosectioning is used to produce sections for imaging applications, and thicker sectioning results in tissue fracture. Drawing on past literature^23^, we carried out a series of experiments testing several sectioning thicknesses and punch diameters followed by nuclei isolation and debris-cleanup on flash-frozen, embedded mouse brain microbiopsy punches. We found that holding cryo-chamber and chuck temperatures at −11°C improves flexibility of the fragile fresh-frozen tissue while maintaining adherence of embedded tissue to the sample mount, thus allowing for uninterrupted sectioning of alternating 100-300 μm sections for punching, and paired 20 μm sections for histology (Figure 1a). This approach facilitates acquisition of both sections for microbiopsy punching and paired sections compatible with IHC staining and high-resolution microscopy. Cryopreservation of 100-300 μm/20 μm slide decks at −80°C allows for long-term sample storage and the ability to test hypotheses by staining after analysis of the spatially resolved chromatin accessibility profiles; however, we note that sections stored for ~3 months result in an overall loss of quality in transcription start site (TSS) enrichment.

Microbiopsy punching of 100-300 μm sections performed within a cooled chamber (Methods) allows for isolation of microscopic pellets of nuclei that readily dissociate in nuclear isolation buffer after mechanical dissociation by trituration. We observed minimal loss after pelleting and washing nuclei, an important step for the removal of mitochondria, which can deplete the available pool of transposase because of the high transposition efficiency into mitochondrial DNA^24^. Nuclei isolation, as measured by nuclei per cubic micron, was more efficient for volumetrically smaller punches (Figure 1b). This implies that smaller punches dissociate more readily because of a higher surface area to volume ratio, thus higher resolution punches yield more nuclei, respective of volume.

Next, we applied these techniques to perform sciMAP-ATAC, where we tested four methods of punch dissociation (Methods). We utilized a workflow similar to our established sci-ATAC-seq method, with each indexed transposition reaction performed on an individual punch, for a total of 384 transposition reactions, performed in four 96 well plates. Reactions were pooled and indexed nuclei were distributed via Fluorescence Assisted Nuclei Sorting (FANS) to wells of 4 new 96-well plates for indexed real-time PCR, followed by pooling and sequencing. The resulting library produced 8,011 cells passing filters, for an estimated doublet rate of 2.5% based on the total indexing space of 384 × 384 (Methods), and a mean of 12,052 passing reads per cell (unique reads, aligned to autosomes or X chromosome at q10 or higher; Extended Data Figure 2a) at the depth sequenced and potential to reach 23,830 mean passing reads per cell with additional sequencing (Methods). This is comparable to the mean passing reads per cell from the whole brain sci-ATAC-seq library at 11,987 (projected mean passing reads of 24,672 and 32,029 for fresh and frozen preparations, respectively; Figure 1c, Extended Data Figure 2a). We observed a mean of 112 passing cells per punch. This could be increased if additional PCR plates were sorted, as the pool of indexed nuclei were not depleted during FANS. A comparison between the four dissociation methods enabled us to identify an optimal means of punch processing that produced the highest cell counts per punch with high-quality cell profiles (Methods; Extended Data Figures 1b and 2a), which was used for all subsequent experiments. Across all sciMAP-ATAC datasets produced in this study on healthy mouse brain tissue, we achieve a TSS enrichment of 14.73, within the ‘acceptable range’ prescribed by ENCODE (10-15, mm10 RefSeq annotation) and just shy of ‘ideal’ (>15). This is substantially below that of our sci-ATAC-seq preparation, with a TSS enrichment of 31.25; however, we note that an enrichment of more than double the ‘ideal’ standard is exceptionally high (Methods, Figure 1d). In line with the lower TSS enrichment in sci-MAP-ATAC, we also observed a reduction in the fraction of reads present in a mouse reference peak set (FRiS; Methods), with a mean ranging from 0.83 to 0.87, compared to 0.91 and 0.93 for sci-ATAC-seq (Extended Data Figure 2b). Finally, we performed an integrated analysis across these preparations that revealed negligible batch effects (Figure 1e, Extended Data Figures 3a and 3b). We observed a single exception in the form of a population of cells present only in the non-spatial dataset which, upon inspection, were determined to be spinal cord derived interneurons (Extended Data Figure 3c and 3d) and not present in coronal sections that were used in spatial experiments. Taken together, with improvements and validation on sample preparation, cryosectioning, nuclei isolation and the general sci-ATAC-seq protocol, we generated a robust method to obtain spatial information that we sought to test in a complex system.

### sciMAP-ATAC in the adult mouse somatosensory cortex

To establish the ability of sciMAP-ATAC to characterize single cells within a spatially organized tissue, we applied the technique to resolve murine cortical lamination within the primary somatosensory cortex (SSp). We harvested intact whole brain tissue from three wild-type C57/Bl6J adult male mice, flash-froze the tissue, and prepared whole brain slide decks of 200 μm microbiopsy slides each interspersed with three 20 μm histological slides. To orient sections to intact mouse brain, and to establish the quality of histological section prepared according to the sciMAP-ATAC protocol, we stained nuclei using DAPI and immunohistochemically stained for Satb2 to resolve cortical layers (Methods, Figure 2a). DAPI imaging was then matched to the adult mouse Allen Brain Reference Atlas^25^, which enabled determination of the SSp location within adjacent sections for punch acquisition. Satb2 imaging demonstrated the quality of histological sections, across diverse fixation protocols (4% PFA post-fixation for 10 minutes and 70% ethanol post-fixation for 30 seconds) and generated a high signal-to-noise ratio canonical for Satb2 IHC staining^26^ (Figure 2b). Microbiopsy punches were then taken from three regions: i) outer (L2-4) SSp cortical layers, ii) inner (L5,6) SSp cortical layers and iii) throughout the striatum. The striatum is rich in glia and is absent of cortical glutamatergic neurons and cortical lamination. Therefore, the striatum punches served as a negative control for these features and also bolstered single-cell glial cell type identification. In total, 96 individual tissue punches were obtained, split evenly between the three categories over eight coronal sections spanning the SSp (Figure 2a). After nuclei isolation, each well of the plate containing a single punch was split across four total 96-well plates for subsequent indexed transposition, providing four tagmentation technical replicates for each punch. Post-transposition, nuclei were pooled and distributed to two 96-well PCR plates for the second tier of indexing and then sequenced using previously described protocols^22^.

**Figure 2.**
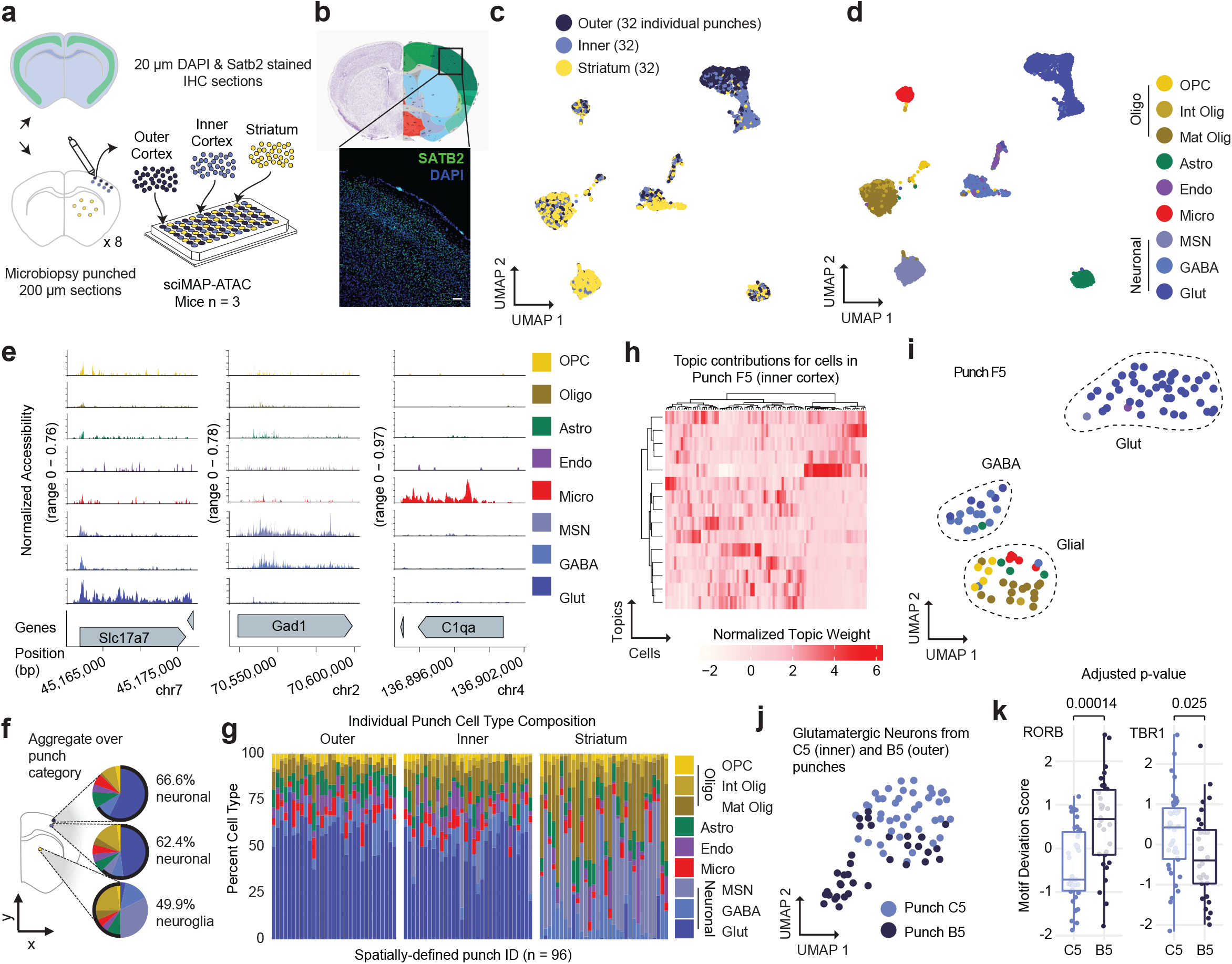
sciMAP-ATAC reveals spatially distinct cell type composition in the mouse somatosensory cortex. **a.** Experiment schematic of sciMAP-ATAC in the mouse somatosensory cortex. **b.** DAPI and SATB2 staining of SSp cortex from sciMAP-ATAC histological section (scale bar, 50 μm). **c.** UMAP of 7,779 cells colored by punch location category. Each category contains cells from 32 spatially distinct tissue punches. **d.** UMAP as in c. colored by cell type. **e.** ATAC-seq profiles for cells aggregated by cell type for marker genes. **f.** Aggregate cell type composition over punches belonging to the broad region categories. **g.** Cell type composition for each of the 96 individual punches split by broad region category. **h.** Topic weight matrix for cells present only in a single punch (F5, inner cortex punch). **i.** UMAP of cells from punch F5 showing spatially distinct groupings for cell types. Cells are colored by cell identity from panel d. **j.** UMAP of glutamatergic neuron cells from two adjacent punches (C5, inner cortex, and B5, outer cortex) after topic modeling on the isolated cell profiles. **k.** Transcription factor motif enrichments for glutamatergic cells from adjacent punches shown in j. Two-sided Mann–Whitney *U* test with Bonferroni-Holm correction. Center value represents median.

We processed the raw sequence data as previously described^22^, which resulted in 7,779 cells passing quality filters (estimated doublet rate of 4.9%; Methods). Our mean passing reads per cell was 17,388, with a projected total passing mean reads per cell of 37,079 (Methods) and a TSS enrichment ranging from 13.74 to 15.26 (Extended Data Figures 2a and 2c). A median of 81 single-cell profiles was obtained per punch, with little bias for punch target region or section (Extended Data Figure 2c). Subsequent peak calling, topic modeling, and dimensionality reduction (Methods) revealed cell groupings that were either mixed between the three regional categories or highly enriched for cells derived from the cortex, which was further divided by outer versus inner punch location (Figure 2c, Extended Data Figures 3e and 3f; Extended Data File 2). Overlay of spatial data on the UMAP projection fits with our expectation that glutamatergic (excitatory) neurons are cortex exclusive, displaying an absence of punch-to-punch crosstalk or contamination. Additionally, these cells were integrated with prior sciMAP-ATAC, and sci-ATAC-seq experiments where excitatory neuron clusters were also dominated by cortex-derived punches, with a shared spatial bias between upper and lower punch positions. This demonstrates that spatial datasets can be integrated with non-spatial datasets to provide additional spatial information to those datasets using label transfer or other analysis techniques (Extended Data Figures 3a and 3b).

We performed graph-based clustering to produce eleven clusters over eight broad cell type groups corresponding to glutamatergic neurons, GABAergic (inhibitory) neurons, GABAergic medium spiny neurons (MSNs; also referred to as spiny projection neurons), oligodendrocyte precursor cells (OPCs), newly formed oligodendrocytes, mature oligodendrocytes, astrocytes, microglia, and endothelial cells based on the chromatin accessibility signature of regulatory elements proximal to marker genes (Methods; Figures 2d and 2e; Extended Data File 1). GABAergic neurons co-cluster independent of punch location. Instead, these neurons subdivide into non-layer-specific cell sub-types in the cortex and striatum. However, glutamatergic neurons separate along the dorsal-ventral axis. This recapitulates known neuronal cell state biology, where glutamatergic pyramidal neurons express cortical layer(s) specific markers that define the spatially defined cortical layers. Within the SSp derived cells, we observed 66.6, 62.4, and 49.9 percent of cells corresponding to neurons in the inner cortex, outer cortex, and striatum, respectively. These equate to glia to neuron ratios (GNRs) of 0.50, 0.60, and 1.00 from inner cortex, outer cortex, and striatum, respectively, which correspond to previously reported mouse cerebral cortex and striatum GNRs of 0.66 and 0.97, respectively^27^. In addition to coarse cell type characterization across the major punch categories, we also determined cell type composition for each individual spatially resolved punch (Figure 2g). For cortical punches, little variance was observed within the outer and inner punch categories; however, we did observe increased variability in the proportion of MSNs in the striatum punches, ranging between 2.78 and 72.64 percent, suggesting a non-even distribution of these cells, which is confirmed by MSN cell-type marker, DRD1, IHC staining in adult C57BL/6j striatum (Allen Mouse Brain Atlas).

We next characterized the single-cell ATAC profiles produced from a single tissue punch. We isolated cell profiles that were from punch F5 (n=90 cells), an inner cortex punch, and performed the same analysis as above using the set of peaks called on the full dataset. This produced a set of topic weights that contained a clear structure and were associated with specific cell types (Figure 2h). This was also clear in the UMAP projection, with three primary clusters of cells identified (Figure 2i, Extended Data Figure 3g). Each of these groups was dominated by one cell type, including glutamatergic neurons and GABAergic neurons, with the third group comprised predominantly of glial cell types. We then took the examination of this individual punch further by performing all aspects of the analysis, including peak calling, on only the cell profiles present in punch F5. From those 90 cells, we were able to call 8,460 peaks which were sufficient to perform topic modeling and UMAP visualization, and identify two distinct clusters: one comprised of glutamatergic neurons, and the second containing all other cell types, based on the cell type identities established in the analysis of the full dataset (Extended Data Figures 3h and 3i). A comparison of global motif enrichment between the two clusters revealed elevated Neurod6 and Tbr1 and depleted Sox9 motif accessibility in the cluster comprised of glutamatergic neurons, suggesting very coarse cell type class assignment can be performed on data from a single punch analyzed in isolation (Extended Data Figure 3j). Further resolution of cell types on such a small number of cells, especially without leveraging larger peak sets, is not likely feasible simply due to the low abundance of certain cell types - for example, there was only one endothelial cell present in punch F5. However, it is unlikely that individual punches would be profiled alone in an experiment and the throughput provided in sciMAP-ATAC enables identification of low-abundance cell types in the aggregate dataset, which can be used when performing analysis on individual punch positions.

Finally, we explored whether we could identify and characterize spatially distinct chromatin properties from a single cell type present within two adjacent punches. We isolated cells that were identified as glutamatergic neurons in two punches, C5 (inner cortex) and B5 (outer cortex), that were immediately adjacent with 83 and 65 total cells, and 42 and 35 glutamatergic cells, respectively. Similar to the single punch analysis, we produced a counts matrix including only these cells and used the full set of peaks to perform topic analysis and visualization using UMAP, which showed clear separation between the two locations (Figure 2j). We then assessed global motif accessibility, which revealed clear enrichment for motifs associated with upper or lower cortical layers, including RORB, enriched in the outer cortex, and TBR1, enriched in the inner cortex (Figure 2k).

### Spatial trajectories of single-cell ATAC-seq in the human cortex

With the ability to probe spatial single-cell chromatin accessibility established in the mouse cortical lamination experiment, we next deployed sciMAP-ATAC on human brain tissue to profile lamination in the adult primary visual cortex (VISp) using an equivalent resolution of 215 cubic microns. Samples of human VISp tissue were obtained from an adult (60 yr., male) with no known neurodegenerative disorders at 5.5 hours postmortem. Samples were oriented and flash frozen in TFM prior to storage at −80°C. The sample was cryosectioned using the same alternating thick (200 μm) and thin (20 μm) pattern as previously described. We designed and implemented a 250 μm diameter punch schematic across three adjacent 200 μm sections to produce twenty-one distinct trajectories comprised of eight punches spanning the cortex, with an additional twenty punches distributed in the subcortical white matter for a total of 188 spatially mapped tissue punches (Figure 3a). In total, 4,547 cells passed quality filters with a mean of 30,212 reads per cell (estimated mean of 98,274 passing reads per cell with additional sequencing; Methods, Extended Data Figures 2a and 4a), a mean TSS enrichment of 15.80 – more than twice the ‘ideal’ ENCODE standard for bulk ATAC-seq datasets (>7, GRCh38 RefSeq annotation), and a FRiS of 0.45 using a human reference dataset^28^ (Methods, Extended Data Figures 2b and 2d).

**Figure 3.**
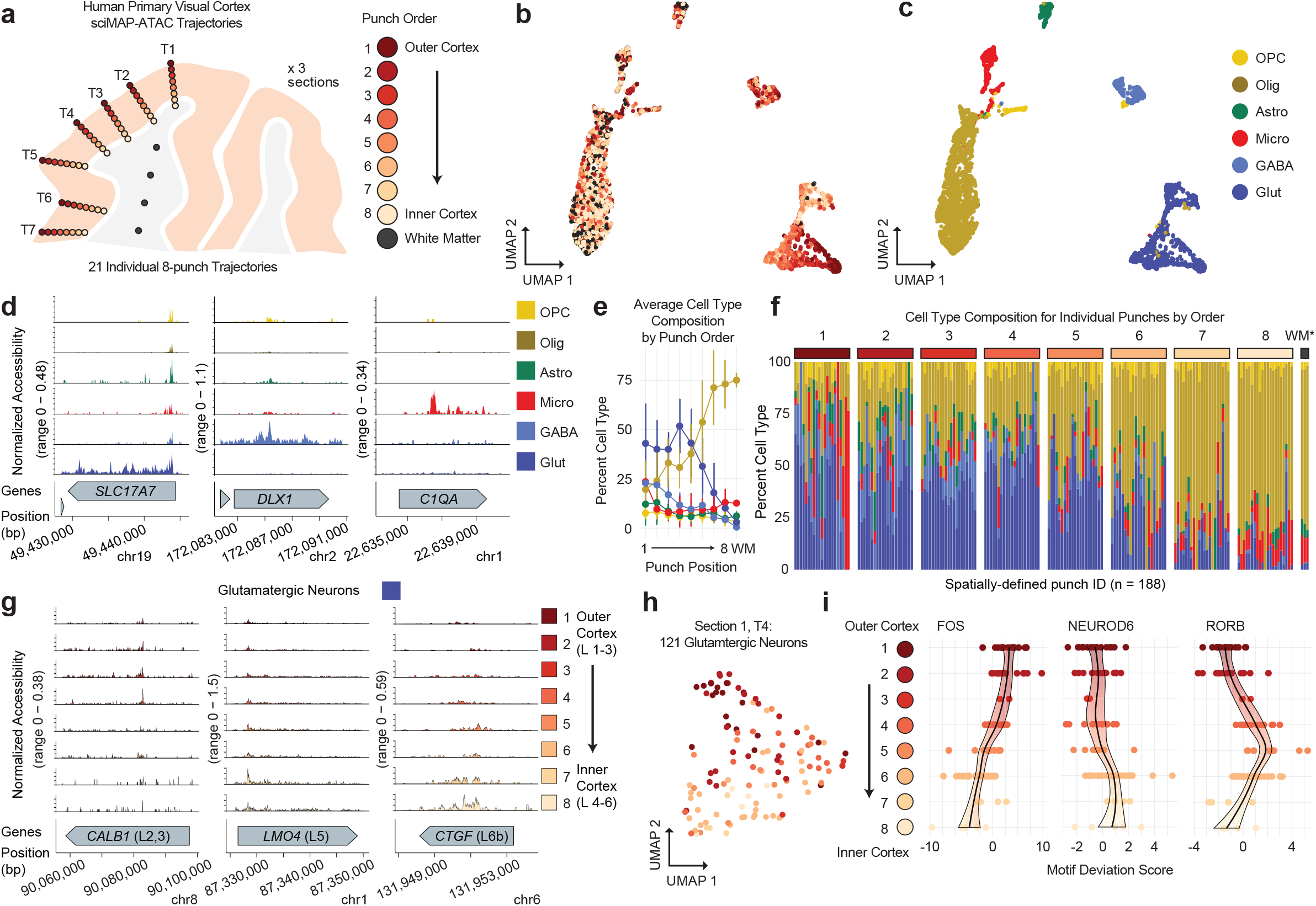
sciMAP-ATAC trajectories through the human primary visual cortex. **a.** sciMAP-ATAC punching schematic showing one of three adjacent sections. A total of 21 eight-punch trajectories spanning the cortex were produced. **b.** UMAP of cells colored by position within their respective trajectory. **c.** UMAP as in b. colored by cell type identity. **d.** ATAC-seq profiles for cells aggregated by cell type for marker genes. **e.** Aggregate cell type composition across the 21 trajectories. Colors are the same as in d. **f.** Cell type composition for each of the 188 individual punches split by trajectory position. Punches from the WM indicated by an asterisk are aggregated by section. Colors are the same as in d. **g.** ATAC-seq profiles for glutamatergic neurons along trajectory positions for layer-specific marker genes CALB1 (layers 2 and 3), LMO4 (layer 5) and CTGF (layer 6b). **h.** UMAP of glutamatergic neurons from a single trajectory (Section 1, Trajectory 4: T1.4) after topic modeling on the isolated cells. Cells are colored by position along the trajectory. **i.** DNA binding motif enrichment for layer-specific factors for Trajectory 1.4 shown in panel h. with cells split by their positions along the trajectory.

Cell profiles were generated as described in prior experiments, which resulted in six distinct clusters representing the major cell types. Similar to the murine cortex, glutamatergic neurons exhibited the most distinct spatial patterning with a clear gradient spanning cortical trajectories (Figures 3b-3d). Each of the 21 individual trajectories through the cortex produced similar distributions of cells through UMAP projections with a lack of glutamatergic neurons present in the punches obtained from subcortical white matter (Extended Data File 2). Average cell type composition along these trajectories revealed the expected pattern of an increased proportion of oligodendrocytes and decreased glutamatergic neuron abundance as the trajectory approached or entered the subcortical white matter region (Figure 3e). Individual punches largely matched the corresponding average position profile (1-8, WM), with higher variability at the first punch where some trajectories overlapped the pial surface of the cortex (Figure 3f). Using our cell type assignments, we next isolated glutamatergic neurons and split them by position along their respective trajectories. We examined ATAC signal at layer-specific marker genes broken down by each spatially distinct category, which revealed increased accessibility at genes associated with outer cortical layers within the outer cortical punches and vice versa (Figure 3g).

We next selected all cells from the centermost trajectory of section 1 (T1.4, n=358 cells) and performed an isolated analysis using peaks called on the full dataset for topic analysis, cluster identification, and visualization with UMAP (Extended Data Figures 4b-4e). Clear separation was observed between major cell types across six clusters, with two distinct clusters of oligodendrocytes, two clusters of glutamatergic neurons, one cluster comprised of GABAergic neurons, and finally, a cluster made up of all other cell types (astrocytes, endothelial and OPCs). When performing the analysis in isolation using only T1.4 cells for peak calling, we identified 16,493 peaks that were used for subsequent analysis to produce four clusters with notably less cell type separation than when leveraging the set of peaks from the full dataset (Extended Data Figure 4f-4g). The first cluster was comprised of both glutamatergic and GABAergic neurons, the second was primarily oligodendrocytes, the third included oligodendrocytes as well as the majority of cells from all other non-neuronal cell types, with the fourth cluster comprised of only a handful of cells with no dominant cell type. In line with the previous assessment of a single punch from the mouse SSp, cell type separation can be distinct for major cell types when leveraging larger peak sets than the limited number that can be called on small cell count datasets. This supports the assertion that computational improvements to enable peak calling on low cell count datasets can substantially boost analytical power^29^.

Finally, we isolated only cells determined to be glutamatergic neurons based on the full dataset cell type assignment within Trajectory 1.4 (n=121 cells). We assessed these cells again using the full peak set through the same analysis workflow as previously described. As in the UMAP projections on cells from the full experiment, these cells were positioned along a gradient that reflected their position along the trajectory (Figure 3h). We then assessed the global accessibility of DNA binding motifs that captured spatially distinct enrichments through the trajectory reflecting the expected pattern of transcription factor activities through cortical layers (Figure 3i). This included enrichment for FOS motif accessibility in the outer cortical layers, slightly increased accessibility for NEUROD6 toward the inner cortex, and increased accessibility for RORB motifs in punches 4-6 along the trajectory, corresponding to canonical cortical layer 4 RORB expression. Taken together, sciMAP-ATAC is capable of producing high-quality single-cell ATAC-seq profiles from postmortem human tissue with a spatial resolution capable of identifying the major components of cortical lamination, with the capability to characterize a single spatial trajectory through the cortex.

### sciMAP-ATAC in cerebral ischemia reveals spatially progressive chromatin features

Beyond the characterization of architecture in structured organs, spatially resolved techniques are particularly well-suited to investigate injury or disease pathology that includes a spatially graded response. Cerebral ischemia produces a complex spatially progressive phenotype with extensive tissue alterations and shifts in cell type abundance and epigenetic states^30–35^. Cerebral ischemic infarction, or cell death as a result of impaired blood flow in the brain, is followed by gliosis, a process in which glia in the surrounding tissue enter reactive states that are potentially aimed at restoring tissue homeostatis, but can involve the loss of normal function (or adopt of damaging function) and form a glial scar. Many components involved in the ischemic cascade are well studied, including factors that promote post-ischemic inflammation (e.g. IRF1, NF-kB, ATF2, STAT3, EGR1, and CEBPB) and prevent post-ischemic inflammation and neuronal damage (e.g. HIF-1, CREB, C-FOS, PPARα, PPARγ, and P53)^36^. Reactive gliosis can be characterized by increased *Gfap* expression in astrocytes and increased *Iba1* in microglia. Myelination depletion is a hallmark of cerebral ischemic injury, due to acute oligodendrocyte cell death and impaired OPC differentiation^37,38^. Far less is known, however, about glial cell state transitions in the area surrounding focal ischemic infarction in the brain. We reasoned that our sciMAP-ATAC technology could reveal, with cell type and spatial specificity, the epigenetic alterations that occur to accompany and/or drive the ischemic cascade and post-ischemic pathology.

To accomplish this, we used a transient middle cerebral artery occlusion (MCAO) mouse model of ischemic injury with reperfusion (Figure 4a). Each ischemic (n=2 animals) and naïve (n=3) brain was flash frozen 3 days after surgery, embedded in TFM, sectioned, alternating between 200 μm for sciMAP-ATAC and 20 μm for IHC for IBA1 (microglia), GFAP (astrocytes, Figure 4b), and counterstained using DAPI. We used these images to define the infarct area by lack of autofluorescence and absence of GFAP-positive astrocytes while being surrounded by reactive astrocytes exhibiting increased GFAP signal at the infarct border (Extended Data Figure 5a). We next defined two axes for targeting the sciMAP-ATAC punches, the first progressing from the pial surface of the cortex to the striatum, all within the infarct core (punch position axis 1-4), and the second progressing from the infarct core toward the infarct border (punch position axis 5-8). GFAP staining intensity as measured by IHC was absent in the infarct core (punch positions 5-7) but increased at the infarct border in punch position 8, recapitulating known features of glial scar formation surrounding the infarct area. We then performed sciMAP-ATAC on the 200 μm sections along each axis to produce 5,081 cells with a mean passing reads per cell of 33,832 (estimated mean passing reads per cell of 225,670 with further sequencing) and a mean of 26.6 high-quality cell profiles per punch (Extended Data Figures 2a, 2e, and 5b). TSS enrichment for this preparation was notably lower than previous preparations ranging from 5.05 (stroke hemisphere) to 7.50 (naïve brain), which we suspect is due to several factors (Extended Data Figure 2e). The first is that the stroke hemisphere contained many dead or dying cells that exhibit reduced ATAC signal, which we describe in more detail below, and the second is that these sections were stored for >3 months prior to sciMAP-ATAC processing, suggesting that long-term storage of sections may result in a reduction in data quality. Despite the reduced TSS enrichment and comparably lower FRiS (0.79-0.82; Extended Data Figure 2b), we called 140,772 accessible genomic loci that were used in subsequent analysis.

**Figure 4.**
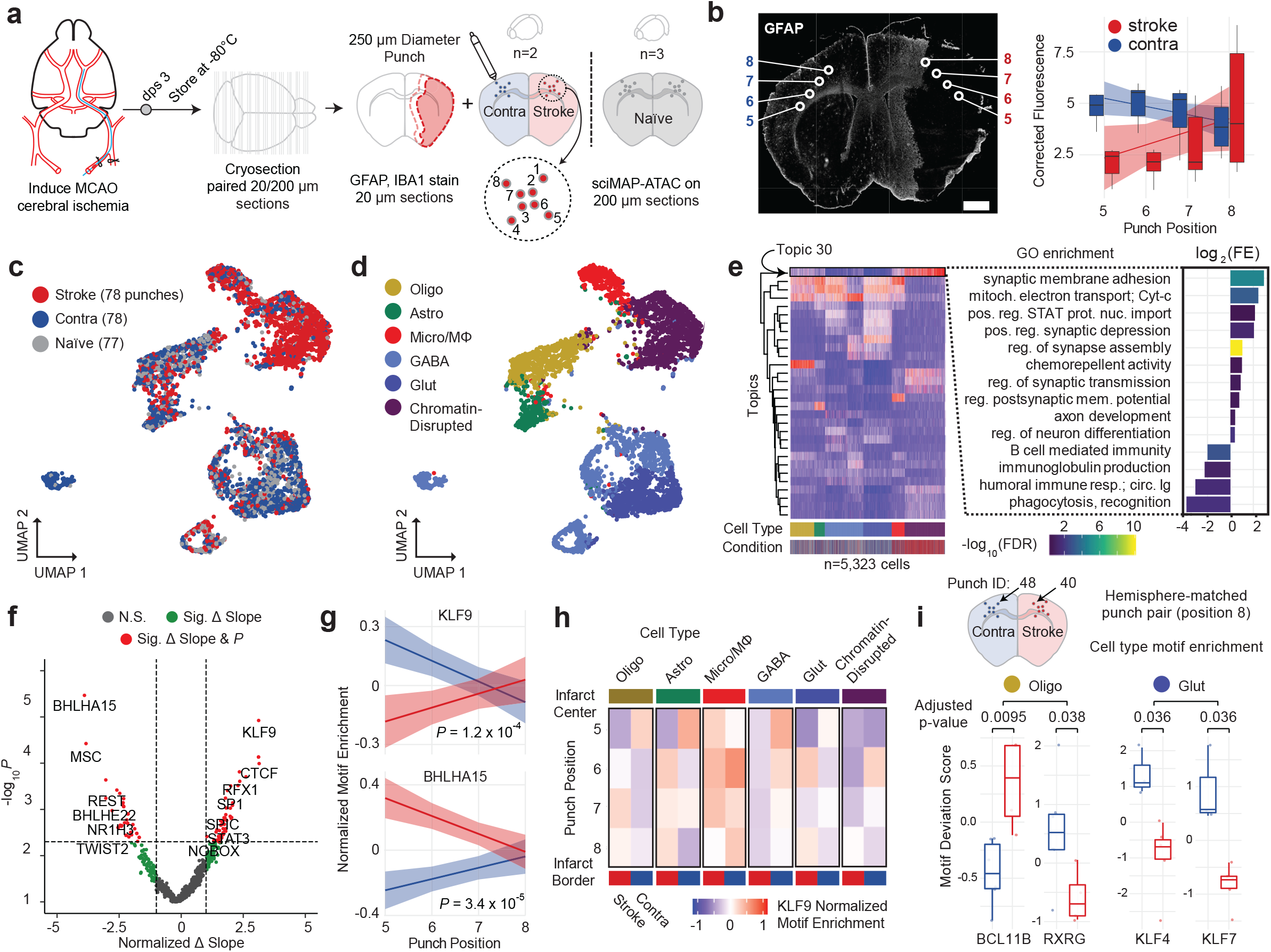
sciMAP-ATAC reveals spatially progressive epigenetic disruption in ischemic injury. **a.** Experimental design using a mouse MCAO model of ischemic injury. Mice were sacrificed three days post-surgery and brains flash frozen in TFM. Alternating thin (20 μm) and thick (200 μm) sections were processed using IHC to define infarction (red outline) and peri-infarct area (pink outline) and sciMAP-ATAC punching schematic, respectively. **b.** GFAP IHC of a 20 μm coronal section of an ischemic mouse brain. Punch positions along the 5-8 axis (core-to-border) are indicated. Background corrected GFAP fluorescence along the 5-8 axis is shown to the right for stroke and contralateral hemispheres. Line shows linear fit ± s.e. (Scale bar, 1 mm) **c.** UMAP of cells colored by the three conditions. **d.** UMAP as in (c) colored by clusters assigned to cell types. **e.** Cell × Topic matrix colored by topic weights, and annotated by cell types and conditions (bottom) reveals substantially divergent topic weighting in cells from the stroke punches (left). Topic 30, enriched specifically in the stroke cells belonging to the chromatin-disrupted cluster has peaks enriched for ontologies associated with ischemic injury with reperfusion. Colored by −Log_10_ false discovery rate (FDR) Q-value, height by log_2_ Fold Enrichment. **f.** Volcano plot of Z-scored TF enrichment slope change across punches 5-8 (Δslope = slope_stroke_−slope_contralateral_) by −log10 *P* value of the two-way ANOVA from the interaction of TF motif enrichment per punch by condition (stroke, contralateral). **g.** Top hits for significantly different changes in TF motif enrichment over space as compared between stroke and contralateral; Klf9 (top) and Bhlha15 (bottom). −Log10 *P* value of the two-way ANOVA from the interaction of TF motif enrichment per punch by condition (stroke, contralateral). Line shows linear fit ± s.e. **h.** KLF9 TF motif enrichment over space reveals cell type contribution to Klf9 enrichment from infarct core to peri-infarct area. **i.** Comparison of TF motif enrichment at the infarct border (punch position 8) between stroke (punch 40) and contralateral (punch 48) single-cell profiles. Oligodendrocyte TF motif enrichment shown for BCL11B and RXRG. Glutamatergic neuron TF motif enrichment shown for KLF4 and KLF7. Two-sided Mann–Whitney *U* test with Bonferroni-Holm correction. Center value represents median.

We performed topic modeling on the cell × peak matrix followed by clustering, cell type identification, and visualization on the cell × topic matrix (Figures 4c and 4d), which revealed comparable cell type proportions across biological samples with exceptions for microglia/macrophages and a chromatin-disrupted cluster that were highly enriched within the infarct. We profiled cell type proportions along both of the axes (Extended Data Figures 5c); however, the pial to striatum axis (punch positions 1-4) in stroke hemisphere samples is completely within the infarct core. In contrast, the infarct core-to-border axis (punch positions 5-8) progresses from the center of the infarct to the glial scar along the infarct border, capturing a transition zone of reactive gliosis, and is the spatial trajectory that we focus on in our subsequent analysis. Along this progression, we found that the stroke hemisphere had diminished neural cell types (depletion of glutamatergic and GABAergic neurons, oligodendrocytes and astrocytes) as well as a progressive increase in cells within a cluster exhibiting globally disrupted chromatin structure up to punch position 7 and a drop at punch position 8 upon entering the infarct border (Extended Data Figure 5d). This state is predominantly characterized by globally increased chromatin accessibility, with a decrease in TSS enrichment, a decrease in FRiS, and an increase in reads falling within distal intergenic regions, which is likely caused by cell death (Extended Data Figures 5e and 5f). In addition to the global effects on chromatin structure, the chromatin-disrupted cell population also showed strong enrichment in one of the topics (Topic 30; Figure 4e, left). A gene ontology (GO) enrichment analysis of the peaks that define Topic 30 revealed that cells within the ischemic hemisphere undergo a chromatin state shift as a result of the ischemic cascade, which leads to enrichment for processes canonically associated with ischemia (Figure 4e, right). Most notably, positive regulation of synaptic membrane adhesion, synaptic depression, assembly, transmission, and membrane potential were all enriched in ischemia-derived cells, indicating that CNS synaptogenesis is upregulated in a subset of cells three days post-ischemia^39,40^. Additionally, while the percentage of microglia increased in the stroke condition (13.2%) as compared to contralateral (6.7%) and naïve (4.3%), depletion of immune response processes (B-cell mediated immunity, humoral immune response mediated by circulating immunoglobulins) were seen in ischemia-derived cells. This recapitulates previous findings that acute ischemic immune response is followed by post-stroke immunodepression and dysregulation^41,42^.

To directly characterize the relationship between space and epigenetic state in cerebral ischemia, we assessed transcription factor (TF) DNA binding motif enrichments for each cell and performed a regression for all cells across the infarct core-to-border axis (punch positions 5-8) in the stroke and contralateral hemispheres. We used the difference between linear model coefficients for paired affected (stroke) and unaffected (contralateral) hemispheres along with the significance of the hemisphere motif enrichment differences to identify putative TFs that undergo spatially progressive regulatory changes (Methods). In total, we identified 56 TF motifs that were significantly altered with a spatial component, many of which have been previously reported as key factors identified in cerebral ischemia (Figure 4f and 4g). KLF9 demonstrated the most significant increase in accessibility with proximity to the peri-infarct area and is a member of the Krupple-like factor (KLF) family. The seventeen KLF family transcription factors are key factors in neuronal development, plasticity, and axon regeneration and are ubiquitously expressed in the CNS. Several KLF family members, namely KLF2, 4, 5, 6, and 11, have been specifically linked to cerebral ischemia pathogenesis^43,44^. Notably, KLF2 and KLF11 have been shown to contribute to the protection of the blood-brain barrier (BBB) in cerebral ischemia^45–47^. However, as DNA binding motifs within the KLF family are similar, members of the KLF family other than KLF9 may be driving this accessibility change. Finally, we assessed the accessibility of individual elements and identified 3,852 peaks that varied significantly through the 5-8 axis of spatial progression (Methods; Extended Data Figure 5g).

We next explored the cell type specificity of the KLF9 motif accessibility changes (Figure 4h). In the stroke hemisphere chromatin-disrupted cell subset, we observed a reduction in KLF9 motif accessibility in all punch positions except punch position 8, at the infarct border, with all cell types other than microglia showing a reduction in accessibility at the center of the infarct core (punch position 5). Uniquely, microglia are largely unaffected and have comparable KLF9 TF binding motif enrichment at the infarct core in comparison to the contralateral hemisphere. In addition to KLF9, we also identified STAT3 as varying significantly over space (Extended Data Figure 5h), which was also an enriched GO term in stroke cells (Figure 4e). STAT3 has been extensively studied in the JAK/STAT3 pathway, which is a key regulator of apoptosis in cerebral ischemia injuries with reperfusion^48^ as well as an initiator of reactive astrogliosis under diverse conditions^49^. Accordingly, we found that STAT3 was largely absent from astrocytes in punches positions 5-7 but was enriched in the reactive astrocytes at the infarct border zone at punch position 8. In contrast, we find that RE1-silencing factor (REST) is significantly elevated at the ischemic core and decreases with proximity to the infarct border. Accordingly, REST has been shown to form a histone deacetylase complex that is a director repressor of SP1 in cerebral ischemia, a TF we identify as varying significantly over space, in the opposite direction of REST^35^ (Extended Data Figure 5i).

We next sought to characterize chromatin accessibility profiles of cells isolated from a single punch at the glial scar (Figure 4i). To do this, we isolated two punches (punch 40 and 48), both originating from the same section (15_SB2), from punch position 8 of the stroke (punch 40) and contralateral hemisphere (punch 48). We processed the cells in isolation as described in prior individual punch analyses, using the peak set from the full experiment. We performed DNA binding motif enrichment analysis across all cells^50^ and then performed cell-type-specific comparisons for a glial (oligodendrocyte) and neuronal (glutamatergic neuron) cell type. In oligodendrocytes, 56 TF motifs were significantly different between the stroke and contralateral hemisphere, many of which (78.6%) corresponded to higher enrichment in stroke as compared to contralateral. Specifically, we found BCL11B (CTIP2), a negative regulator of glial progenitor cell differentiation to be significantly increased at the glial scar around the infarct (Figure 4i, left)^51^. Conversely, we found RXRG, a positive regulator of OPC differentiation, and remyelination, to be significantly depleted (Figure 4i, left)^52^. Together these findings indicate impaired ability of OPCs to differentiate into mature oligodendrocytes at the glial scar. In glutamatergic neurons, we found neuron-associated TFs such as Neurod2 to be significantly depleted in the stroke hemisphere, which corresponds with decreased neuronal cell types at punch position 8 in the stroke hemisphere. In accordance with our infarct core-to-border axis (punch positions 5-8) analysis, we found that seven of the KLF family of TFs (KLF 2-4, 6-8, and 12) were significantly depleted in glutamatergic neurons at the glial scar in the stroke hemisphere (Figure 4i, right; KLF4 and KLF7 shown). Interestingly, previous studies have found that in response to cerebral ischemia, KLF 4, 5, and 6 are induced in astrocytes, while KLF2 is depleted in endothelia and induced in microglia^53^. With these data we identify that motif enrichment for many members of the KLF family not only significantly vary over space across all cell types, we also indicate novel depletion of multiple KLFs specifically in glutamatergic neurons at the ischemic glial scar.

## Discussion

sciMAP-ATAC provides a low-cost, highly scalable, hypothesis-independent approach to acquiring spatially resolved epigenomic single-cell data with the use of immediately available commercial tools. Additionally, sciMAP-ATAC is translatable to any tissue, culture, or model system compatible with cryosectioning. While many methods rely on signal-to-noise optical detection of densely packed molecules and computationally intensive spatial reconstruction, sciMAP-ATAC encodes nuclear localization directly into each library molecule, allowing for rapid subsetting of cells by localization and mapping of cells across vector space in 3D. We demonstrate the use of sciMAP-ATAC to profile the murine somatosensory cortex, as well as multi-punch trajectories through the human primary visual cortex, recapitulating known marker gene progression through cortical layering and cell type composition based on the category and positioning of spatially registered microbiopsy punches. We further show the utility of sciMAP-ATAC to resolve the progressive epigenomic changes in a cerebral ischemia model system, revealing distinct trends in chromatin accessibility, cell type composition, and cell states along the axes of tissue damage and altered morphology. Application of sciMAP-ATAC to other highly structured systems or tissues with a gradient of disease phenotype will be particularly valuable areas for this technology. The primary limitation of sciMAP-ATAC is that punches are currently performed manually and registered with adjacent imaged sections post-punching. This limits the precision of desired punch positions as well as throughput; however, automated processing of tissue sections using robotics^54^ where punch patterns are designed on adjacent imaged sections and registered to the target section will enable high-precision, as well as increased throughput into the range of thousands. Furthermore, as spatial transcriptomic technologies evolve, they may enable the acquisition of chromatin accessibility information; however, substantial technical hurdles must first be overcome, and profiles produced would be in aggregate over the feature size and not necessarily single-cell. Finally, here we applied the sciMAP strategy to assess chromatin accessibility; however, it can, in theory, be applied to any single-cell combinatorial indexing technique to enable spatially registered single-cell genome^19^, transcriptome^55^, chromatin folding^56^, methylation^57^, or multi-omic^58–60^ assays.

---

**Extended Data Figure 1.**
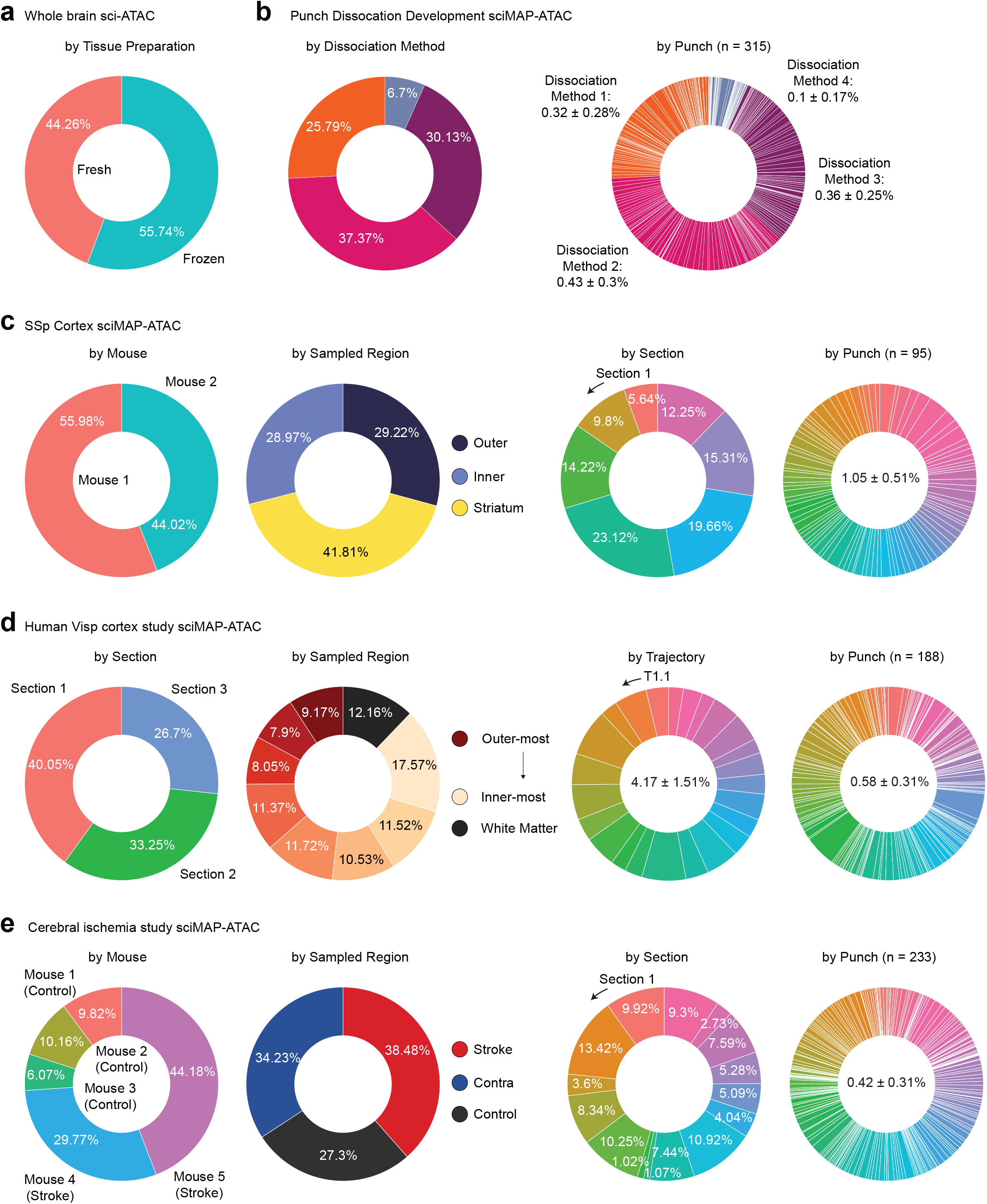
Overview of single-cell ATAC profiles produced across experimental conditions. Quality-passing single-cell ATAC-seq profiles for each experimental condition or spatially resolved punch (for sciMAP-ATAC) as a percentage of the experiment (or average ± standard deviation) for: **a.** sci-ATAC-seq on fresh vs. frozen mouse whole brain hemisphere; **b.** sciMAP-ATAC development across four dissociation methods and individual punches from each dissociation method; **c.** sciMAP-ATAC on mouse SSp by biological replicate, class of sampled region, individual section, and individual punch; **d.** human VISp sciMAP-ATAC by section, class of sampled region, individual trajectory and individual punch; and **e.** sciMAP-ATAC on a mouse model of cerebral ischemia by biological replicate, class of sampled region, section and individual punch.

**Extended Data Figure 2.**
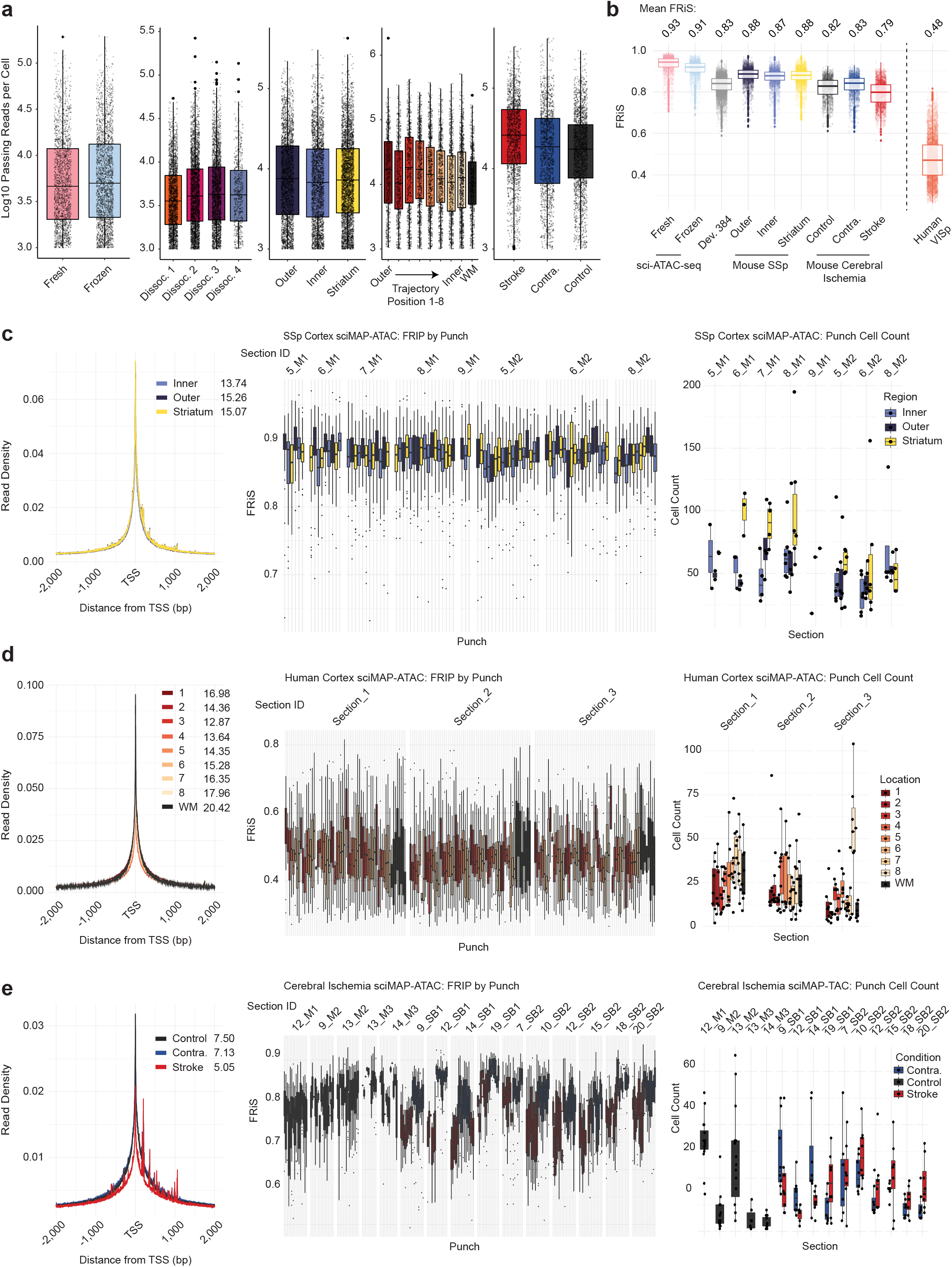
Quality metrics across all experiments. **a.** Log10 passing reads obtained per cell at the depth of sequencing for this study. **b.** The fraction of reads present in a reference set of peaks (FRiS) for each experiment. The master list of peaks for mouse are aggregated from ATAC-seq data produced by the ENCODE project, and for human it is from a single study on DNAse hypersensitivity^28^. **c-e.** Left: aggregate read density centered on TSS sites present in the genome with TSS enrichment values listed by each class calculated using the ENCODE method; middle: FRiS distributions for cells within each punch produced in the experiment split by section; and right: Punch distributions of cell counts for each category within the experiment split by section, for mouse SSp (**c.**), human VISp (**d.**) and mouse cerebral ischemia (**e.**) experiments.

**Extended Data Figure 3.**
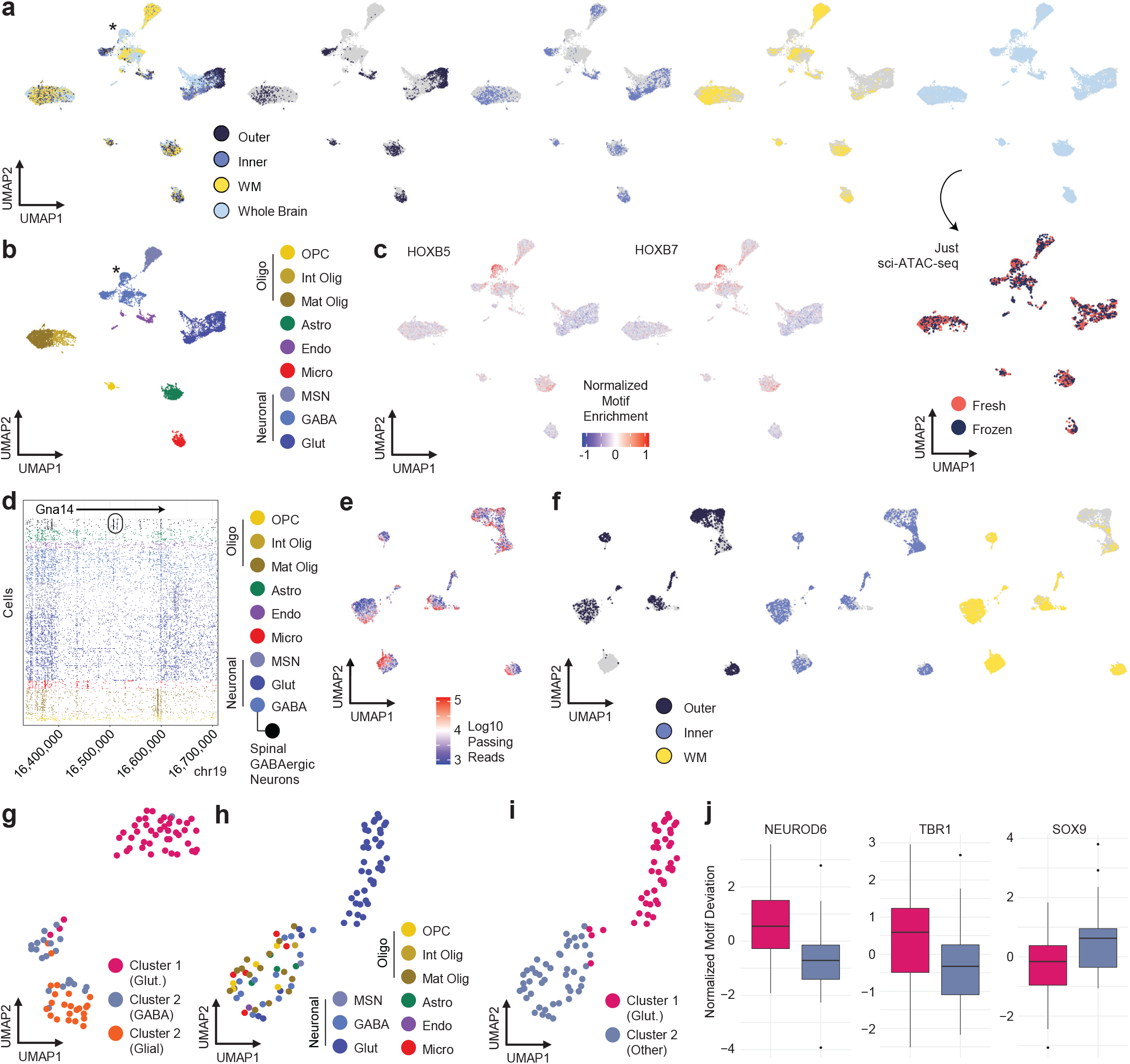
Extended analysis of the mouse somatosensory cortex sciMAP-ATAC dataset. **a.** Integration of all healthy mouse brain sci-ATAC-seq and sciMAP-ATAC datasets visualized in a UMAP. From left to right: all cells colored by either the regional category of punch position for the SSp experiment and then cell from whole brain experiments – asterisk indicates the population of cells only present in the whole brain dataset; cells are grayed out except for those from punches taken from the outer cortex, inner cortex, stratum and then whole brain. Below the whole brain panel, cells from the fresh and frozen sci-ATAC-seq experiments are indicated. **b.** The same integrated UMAP with cells colored by identified cell type. Asterisk indicates the population of GABAergic neurons only present in whole brain datasets that represent spinal cord derived interneurons. **c.** HOXB5 and HOXB7 are two example motifs that exhibit increased accessibility in the spinal cord derived interneuron population. **d.** ATAC reads for cells (rows) are shown for the *Gna14* locus with cells colored by broad cell type. The cluster representing spinal cord derived interneurons is split out and shown in black with the uniquely accessible loci circled. **e.** UMAP of the SSp dataset with cells colored by log10 passing read counts. **f.** UMAP of the SSp dataset with cells grayed out except for each of the three regional punch categories. **g.** UMAP of cells from punch F5 that were processed using peaks called on the full SSp dataset colored by the three clusters that were identified. **h.** UMAP of cells from punch F5 that were processed using peaks called on only those cells colored by their cell type as identified in the full SSp dataset analysis and **i.** colored by the two clusters that were identified. **j.** Motif accessibility for the isolated punch F5 analysis indicating that Cluster 1 is made up of glutamatergic neurons and Cluster 2 is made up of other cell types.

**Extended Data Figure 4.**
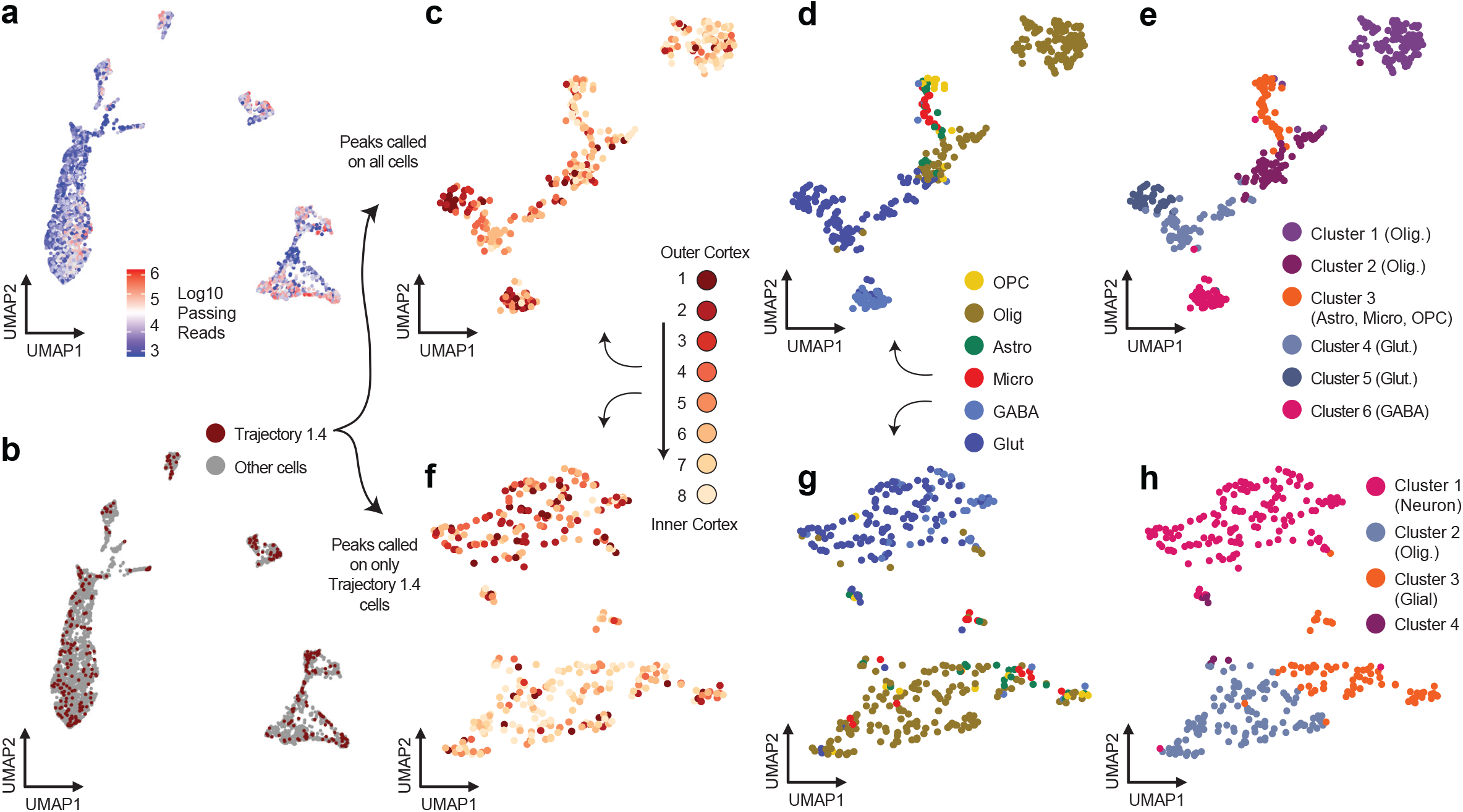
Extended analysis of the human primary visual cortex sciMAP-ATAC dataset. **a.** UMAP of all cells from the experiment colored by log10 passing read counts. **b.** UMAP of the full dataset with all cells grayed out except for those belonging to the middle trajectory of 8 punches on the first section (Trajectory 1.4). **c.** UMAP of cells from Trajectory 1.4 that were processed using peaks from the full VISp dataset colored by the punch position; **d.** the cell type classification as determined from the full dataset; and **e.** the six clusters that were identified. **f.** UMAP of cells from Trajectory 1.4 that were processed using peaks called using only those cells colored by the punch position; **g.** the cell type classification as determined from the full dataset; and **h.** the four clusters that were identified.

**Extended Data Figure 5.**
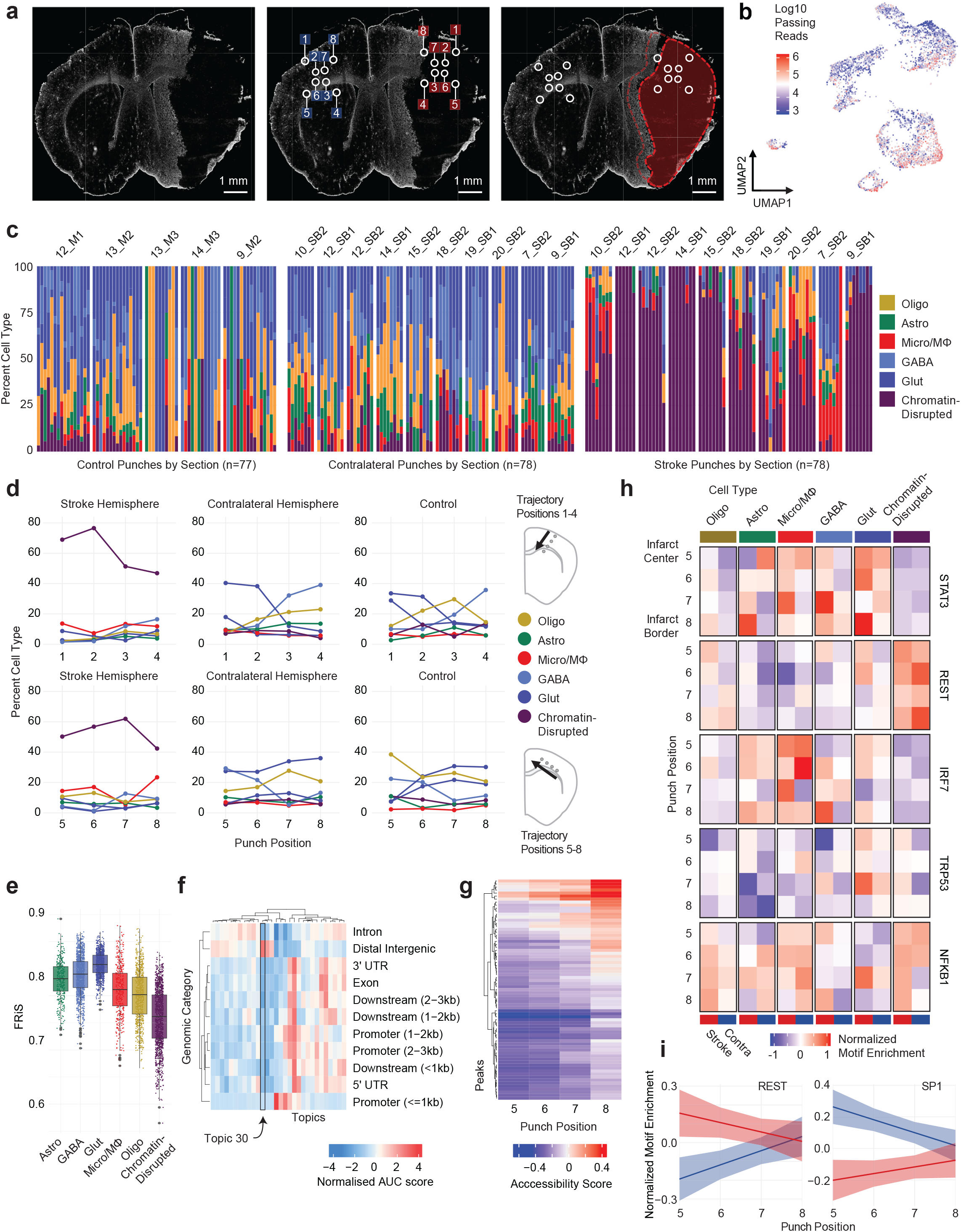
Extended analysis of the cerebral ischemia sciMAP-ATAC dataset. **a.** GFAP staining of an imaging section from a stroke brain (left), with punch positions and labels shown (middle), and punch positions with the stroke region overlaid in red (right). **b.** UMAP of cells from the cerebral ischemia experiment colored by the log10 passing read counts. **c.** Cell type composition for each punch in the experiment grouped by individual section and more broadly by category. **d.** Aggregated cell type composition for the 1-4 axis (top) and 5-8 axis (bottom) split by category of tissue. **e.** FRiS values for cells split by called cell type indicating a substantial decrease in FRiS for the chromatin-disrupted cluster. **f.** Enrichment for topics with respect to genomic category showing that Topic 30, which is elevated in cells within the chromatin-disrupted cluster, is enriched for distal intergenic regions – further supporting a global laxing of chromatin, likely due to cell death. **g.** Regulatory elements that change significantly and uniformly along the 5-8 axis. **h.** Motif enrichment along the 5-8 axis for stroke and contralateral hemispheres split by cell type. **i.** REST and SP1 normalized motif enrichment along the 5-8 axis shows opposite trends between the two factors as well as for each factor between the stroke and contra hemispheres.

**Extended Data File 1 | Marker gene accessibility plots.** Read depth aggregated by identified cell types are plotted for cell type identification marker genes for the mouse SSp, human VISp, and mouse cerebral ischemia sciMAP-ATAC datasets.

**Extended Data File 2 | UMAPs split by each individual punch or trajectory.** UMAPs for the mouse SSp and cerebral ischemia experiment are shown with all cells grayed out except for each indicated punch as well as UMAPs for the human VISp experiment with all cells grayed out except for each individual punch trajectory.

## Acknowledgments

We thank members of the Adey, O’Roak, and Wright labs for their support; Kylee Rosette for assistance with animal husbandry; Eleonora Juarez for discussion on protocol development; Anthony P. Barnes and Brian J. O’Roak for helpful discussions on experimental design; and Dominica Cao and Brooke DeRosa for IHC staining.

## Author Contributions

A.C.A. and C.A.T. conceived of the idea. C.A.T. performed all experiments described with assistance from R.M.M., A.J.F.; F.J.S., K.M.W. and A.M. contributed to experimental design and data interpretation. C.A.T. performed data processing and analysis with assistance from K.A.T., R.M.M., A.N. and E.G.L. W.Z. and H.M. performed stroke surgeries. R.W. identified, isolated, and cryopreserved human primary visual cortex with assistance from C.A.T. C.A.T. and A.C.A. wrote the manuscript with input from all authors.

## Data and Materials Availability

The sciMAP-ATAC protocol is available on Protocols.io (dx.doi.org/10.17504/protocols.io.5r4g58w). Analysis was performed using functions contained within the scitools software (github.com/adeylab/scitools).

## Methods

### Mouse brain and human Visp cortex sample preparation

All animal studies were approved by the Oregon Health and Science University Institutional Animal Care and Use Committee. Male C57Bl/6J mice aged 8 weeks were purchased from Jackson Laboratories for the mouse whole brain sciATAC, punch dissociation development sciMAP-ATAC, and mouse SSp cortex sciMAP-ATAC experiments. Animals were sacrificed by carbon dioxide primary euthanasia and cervical dislocation secondary euthanasia. Animals were immediately decapitated, intact brain tissue was harvested, washed in ice-cold phosphate-buffered saline (PBS; pH 7.4), submerged in TFM (Cat. TFM-C) within a disposable embedding mold (Cat. EMS 70183). Human Visp cortex samples were provided by the Oregon Brain Bank 5.5 hours post-mortem and were submerged in TFM. Embedded mouse whole brain and human Visp cortex samples were flash-frozen in liquid nitrogen cooled isopentane by lowering the sample into the isopentane bath without submerging within 5 minutes of embedding. Samples were immediately transferred to dry ice, paraffin wrapped to delay sample dehydration, and stored in an air-tight container at −80 °C.

### Mouse cerebral ischemia model

Two C57BL/6 9-week-old (P63) female mice were placed under isoflurane anesthesia (5% induction, 1.5% maintenance) in 30% oxygen-enriched air. Body temperature was maintained at 37 ± 0.5°C throughout the procedure. Middle cerebral artery (MCA) occlusion was performed using a previously described method by Longa *et al.* with slight modifications^61^. Briefly, a laser Doppler flowmeter (Moore Instruments) probe was affixed over the right parietal bone overlying the MCA territory to monitor changes in cerebral blood flow. A midline incision was made, the right common carotid artery (CCA) bifurcation was exposed by gentle dissection, and the external carotid artery (ECA) was permanently ligated distal to the occipital artery using electrocautery, such that a short ECA stump remained attached to the bifurcation. The right CCA and internal carotid arteries (ICA) were temporarily closed with reversible slip knots before an arteriotomy was made in the ECA stump. A silicone-coated 6.0 nylon monofilament was inserted into the ICA via the arteriotomy and gently advanced to the ICA/MCA bifurcation to occlude CBF to the MCA territory and confirmed by a laser Doppler signal drop of <30% of baseline. After 60 minutes occlusion, the filament was gently retracted, the ECA permanently ligated, the slip knot of the CCA removed, and the incision sites sutured closed. The mice exposed to MCAO were euthanized three days after the MCAO procedure, intact brain tissue harvested, washed in ice-cold PBS (pH 7.4), submerged in TFM and flash-frozen in liquid nitrogen cooled isopentane. Samples were paraffin wrapped and stored at −80 °C and intact embedded whole mouse brains were sectioned at the time of experiment.

### Sample sectioning

All embedded samples were sectioned in a cryostat (Leica CM3050) at −11°C chuck and chamber temperature and collected on Superfrost Plus microscope slides (Fisherbrand, Cat. 22-037-246). Sectioning was performed in sets of: one section at 100-300 μm paired with three sections at 20 μm, to generate sets of four slides consisting of microbiopsy (1) and histology (3) sections at one section per slide. Slide boxes were sealed with paraffin to prevent sample dehydration and stored long-term at −80°C.

### Mouse whole brain coronal section immunohistochemistry and mapping

To determine the mouse brain atlas coordinate of each coronal microbiopsy section, the histological section immediately adjacent to each microbiopsy section were fixed in 4% PFA for 10 minutes and counter stained using 300 μM DAPI (Thermo Fisher, Cat. D1306) in 1x (pH 7.4) PBS (Thermo Fisher, Cat. 10010023) for 5 minutes. Slides were rinsed with 1x PBS and mounted in Fluoromount-G (Thermo Fisher, Cat. 00-4958-02). Slides stained for Satb2 were equilibrated to room temperature and circumscribed with a hydrophobic barrier pen (Invignome, Cat. GPF-VPSA-V). Sections were washed twice with PBS for 10 minutes then blocked for 1 hour at room temperature in permeabilization/blocking buffer comprised of PBS with 10% normal goat serum (Jackson ImmunoResearch, Cat. 005-000-121), 1% bovine serum albumin (BSA, Millipore, Cat. 126626), 0.3% Triton X-100 (TX-100, Sigma, Cat. 11332481001), 0.05% Tween-20 (Sigma, Cat. P1379), 0.3 M glycine (Sigma, Cat. G7126) and 0.01% sodium azide (Sigma, Cat. S2002). During the blocking step, the primary antibody rabbit anti-Satb2 (Abcam Cat. ab92446) was diluted 1:1000 in a buffer containing PBS, 2% normal goat serum (NGS), 1% BSA, 0.01% TX-100, 0.05% Tween-20, and 0.01% sodium azide. The diluted primary antibody was applied to sections then incubated overnight at 4°C. The primary antibody was washed from the sections five times with PBS for 5 minutes at room temperature. Secondary antibody AF488 goat anti-rabbit (Thermo Fisher Cat. A32731) was prepared by diluting 1:1000 in the same buffer used to dilute primary antibodies. Sections were incubated with the diluted secondary antibody for 1 hour in the dark at room temperature. Secondary antibodies were washed from the sections three times with PBS for 5 minutes, then nuclei were counterstained with DAPI for 10 minutes at room temperature. After DAPI staining, sections were washed an additional two times then glass coverslips were mounted with ProLong Diamond Anti-Fade Mounting Medium (Thermo Fisher, Cat. P36961). Slides were imaged on a Zeiss ApoTome AxioImager M2 fluorescent upright microscope and processed using Fiji software^62^. Coronal section images were mapped to the Adult Mouse Allen Brain Atlas^25^ according to anatomical regions.

### Mouse cerebral ischemia immunohistochemistry and mapping

One of the histological sections corresponding to each microbiopsy section was stained for GFAP to identify the infarct. Slides were equilibrated to room temperature and circumscribed with a hydrophobic barrier pen. Sections were washed twice with PBS for 10 minutes then blocked for 1 hour at room temperature in permeabilization/blocking buffer comprised of PBS with 10% normal donkey serum, 1% bovine serum albumin, and 0.05% Triton X-100. The sections were next incubated in primary antibody solution comprised of 1:1000 goat anti-GFAP (Abcam, ab53554) and 1:5000 rabbit anti-Iba1 (Fujifilm Wako, NCNP24) diluted in PBS with 1% NGS, 0.1% BSA and 0.005% TX-100 overnight at 4°C. The sections were then washed three times with PBS for 5 minutes each at room temperature and next incubated for 2 hours at room temperature in secondary antibody solution containing 1:500 donkey anti-goat conjugated to Alexa Fluor 488 (Invitrogen) and 1:500 donkey anti-rabbit conjugated to Alexa Fluor 555 (Invitrogen) prepared in the same buffer as the primary antibodies. Following the secondary incubation, sections were washed three times with PBS for 5 minutes each, counterstained with DAPI for 10 minutes, washed an additional two times for 5 minutes each, then coverslipped with Fluoromount-G. Slides were imaged on a Zeiss AxioScan.Z1 Slide Scanner and processed using Fiji software. Coronal cerebral ischemia section images were mapped to the Adult Mouse Allen Brain Atlas^25^ according to anatomical regions using the DAPI channel, as described above.

Immunohistochemistry fluorescence was quantified using ImageJ (v1.52p). Punch positions were mapped to regions of interest (ROIs), along with three negative naïve ROIs for each image. Corrected total fluorescence was calculated as the difference between the integrated density (ROI area * Mean fluorescence) of an ROI for a given punch and the average integrated density of negative naïve ROIs. GFAP corrected total fluorescence was plotted using *ggplot geom_boxplot* and *geom_smooth,* method *lm* using the *ggplot* function in R (v3.2.1).

### Mouse whole brain dissociation and nuclei isolation

To evaluate the effect of flash-freezing on chromatin accessibility in mouse brain tissue, we evaluated single-cell chromatin accessibility profiles from an intact mouse brain in which one hemisphere was flash-frozen as described previously, and one hemisphere remained unfrozen. Both hemispheres were processed in parallel and underwent dissociation and nuclear isolation. Tissue was diced in Nuclear Isolation Buffer (NIB: 10mM Tris HCl, pH 7.5 [Fisher, Cat. T1503 and Fisher, Cat. A144], 10mM NaCl [Fisher, Cat. M-11624], 3mM MgCl_2_ [Sigma, Cat. M8226], 0.1% IGEPAL [v/v; Sigma, I8896], 0.1% Tween-20 [v/v, Sigma, Cat. P7949] and 1x protease inhibitor [Roche, Cat. 11873580001]) in a petri dish on ice using a chilled razor blade. Diced tissue was transferred to 2 mL chilled NIB in a 7 mL Dounce-homogenizer on ice. The tissue was incubated on ice for 5-minutes then homogenized via 10 gentle strokes of the loose pestle (A) on ice, a 5-minute incubation on ice, then 10 gentle strokes of the tight pestle (B) on ice. The homogenate was then strained through a 35 μm strainer and centrifuged at 500 rcf for 10 minutes. Samples were aspirated, resuspended in 5 mL of ice-cold NIB, and nuclei were counted on a hemocytometer. Samples were diluted to 500 nuclei per 1 μL to facilitate tagmentation reaction assembly at approximately 5,000 nuclei per 10 μL of NIB.

### Tissue microbiopsy acquisition and nuclear isolation

Tissue microbiopsies were acquired from 100-300 μm sections. Punches were isolated in four experiments: 1) mouse dissociation development sciMAP-ATAC (384 punches), 2) mouse SSp cortex sciMAP-ATAC (96 punches), 3) mouse cerebral ischemia sciMAP-ATAC (240 punches), and 4) human Visp cortex sciMAP-ATAC (192) (for details refer to Extended Data Figure 1). Microbiopsy coronal sections were acclimated to −20° C in a cryostat (Leica CM3050) and microbiopsy punch tools (EMS, Cat. 57401) were cooled on dry ice prior to punching to prevent warming of tissue. Microbiopsy punches were acquired according to location identified from section atlas mapping, and frozen microbiopsies were deposited directly into 100 μL of ice-cold NIB in a 96-well plate. Punch deposition into each well of the 96-well plate was visually confirmed under a dissecting microscope. To facilitate tissue dissociation and nuclear isolation, 96-well plates of microbiopsy punches were then gently shaken (80 rpm) while covered for 1 hour on ice. We then tested mechanical dissociation by varying the number of triturations performed via multi-channel pipette per well (punch dissociation development sciMAP-ATAC). We found the following averaged metrics across the four dissociation methods: 15 triturations (26 cells per punch, 5,679 unique passing reads per cell, 0.844 FRis), 30 triturations (35 cells per punch, 7,189 unique passing reads per cell, 0.835 FRis), 60 triturations (28 cells per punch, 7,611 unique passing reads per cell, 0.827 FRis), and 100 triturations (8 cells per punch, 7,611 unique passing reads per cell, 0.821 FRis). Given that 60 trituration mechanical dissociation yielded the highest number of cells per punch, with otherwise comparable metrics, we proceeded with 60 triturations for all future experiments. Post-mechanical dissociation, sample plates were then centrifuged at 500 rcf for 10 minutes. While nuclear pellets were not visible, we found that aspiration of 90 μL of supernatant and resuspension in an added 30 μL of NIB results in a final isolated nuclear volume of 40 μL with approximately 15,000 nuclei per well (for microbiopsy punching conditions: 200 μm section, 250 μm diameter microbiopsy punch used in the human VISp and mouse cerebral ischemia preparations). Nuclei were split across four 96-well plates such that nuclei were aliquoted to 10 μL, or approximately 3,750 nuclei per well. This enabled 4 independent indexed transposase complexes to be utilized for each individual punch, or 384 uniquely indexed transposition reactions in one experiment. To calculate the approximate resolution for each preparation, we took the cubed root of the cylindrical volume.

### Location indexing via tagmentation

Transposase catalyzed excision of the chromatin accessible regions via tagmentation results in the addition of unique molecular identifiers (indexes) for each tagmentation reaction. To encode microbiopsy punch location into library molecules, we recorded the corresponding tagmentation well within each 96-well plate to the user-identified microbiopsy punch location. The incorporation of location information is therefore inherently encoded by the first tier of indexing in our established sci-ATAC-seq method. Tagmentation reactions were assembled at 10 μL of isolated nuclei at 500 nuclei per 1 μL, 10 μL 2x tagmentation buffer (Illumina, Cat. FC-121-1031), and 1 μL of 8 μM loaded indexed transposase was added per well (See Picelli et al. for loading protocol)^63^. Each assembled 96-well plate of tagmentation reactions was incubated at 55°C for 15 minutes. For the mouse whole brain sci-ATAC-seq preparation on fresh and frozen tissue as well as the sciMAP-ATAC preparations, four 96-well plates of tagmentation were used (384 uniquely indexed tagmentation reactions). For whole brain sci-ATAC-seq preparation on fresh and frozen tissue experiment, tagmentation wells were pooled separately for fresh and frozen hemisphere samples. For the microbiopsy punch-derived experiments, all reactions were pooled post-tagmentation.

### Combinatorial indexing

To lyse nuclei and release bound transposase, PCR plates are prepared with protease buffer (PB), primers, and sparsely sorted nuclei and then incubated. Post-denaturation, the remaining PCR reagents are added and incorporation of the PCR primers results in incorporation of the secondary index for single combinatorial indexing. For the denaturation step, 96-well PCR plates of 8.5 μL PB (30 mM Tris HCl, pH 7.5, 2 mM EDTA [Ambion, Cat. AM9261, 20 mM KCl [Fisher, Cat. P217 and Fisher, Cat. A144], 0.2% Triton X-100 [v/v], 500 ug/mL serine protease [Fisher, Cat. NC9221823), 1 μL 10 mM indexed i5, and 1uL indexed i7 per well were prepared. Pooled tagmented nuclei were stained by adding 3 μL of DAPI (5mg/mL) per 1 mL of sample. Each sample was then FAN sorted using a Sony SH800 FACS machine at 22 events per well per 96-well Tn5 plate (*e.g.* 88 for 384 indexes) into prepared 96-well plate(s). Event numbers were selected based on the expected success rate of events as actual cells for a given target cell doublet rate (see ‘Doublet rate estimations’ section below). Across the sciMAP-ATAC experiments, four PCR plates (384 uniquely indexed wells) were utilized for the initial punch-derived sci-ATAC-seq preparation from whole brain-derived punches, two PCR plates (192 uniquely indexed wells) were used for the mouse SSp cortex experiment, one full and one partial plate (128 uniquely indexed wells) for the human VISp experiment, 2 plates (192 uniquely indexed wells) for the mouse cerebral ischemia experiment, and finally two PCR plates (192 uniquely indexed wells) were utilized for the non-spatial whole brain sci-ATAC-seq preparation on fresh and frozen tissue. Transposase denaturation was performed by sealing each sorted plate and incubating at 55°C for 15 minutes. Plates were immediately transferred to ice post-incubation and 12 μL of PCR mix (7.5 μL NPM [Illumina Inc. Cat FC-131-1096], 4 μL nuclease-free water, 0.5 μL 100× SYBR Green) was added to each well. For each experiment, plates were then sealed and PCR amplified on a BioRad CFX real-time cycler using the following protocol: 72°C for 5:00, 98°C for 0:30, Cycles of [98°C for 0:10, 63°C for 0:30, 72°C for 1:00, plate read, 72°C for 0:10] for 18-22 cycles. PCR plates were transferred to 4°C once all wells reached mid-exponential amplification on average. Each PCR plate is then pooled at 10 μL per well and DNA libraries are isolated using a QIAquick PCR Purification column. Each pooled PCR plate library is then quantified using a Qubit 2.0 fluorimeter, diluted to 4 ng/μL with nuclease-free water, and quantification of library size performed on Agilent Bioanalyzer using a dsDNA high sensitivity chip. Libraries were then sequenced on a NextSeq™ 500 sequencer (Illumina Inc.) loaded with custom primers and chemistry, as previously described^19^.

### Doublet rate estimations

An important factor in single-cell studies is the expected doublet or collision rate. This manifests in droplet-based platforms as two cells being encapsulated within the same droplet, thus having the same cell barcode for their genomic information. This is tunable by the number of cells or nuclei loaded onto the instrument, with typical doublet rates targeted to be at or below 5%. This is also true for combinatorial indexing workflows, where doublets are present in the form of two cells or nuclei with the same level 1 index – which is the transposase index for ATAC – that end up in the same level 2 indexing well (i.e., the PCR well). This results in an identical pair of indexes for the two cells. This rate, like with droplet methods, is also tunable by altering the number of indexed cells or nuclei that are deposited into each well, with a typical experiment targeting at or below a 5% doublet rate. This rate is approximated by leveraging the “birthday problem” formulation in statistics, where the transposase index space (days in the year) and number of indexed nuclei per well (number of people at each table) are taken into account. These predictions assume that there is complete mixing of nuclei prior to distribution and that the distribution is unbiased, which are reasonable given the single nuclei suspension and use of flow sorting for the distribution process, and hold up when compared to empirical data produced by multi-species cell mixing experiments^19,57,64^ (i.e. barnyard experiments, typically mixing human and mouse cells). However, in the case of sciMAP-ATAC, nuclei are directly isolated and then indexed within the same well, making a true barnyard experiment not feasible. Any experiment that would use tissue punches from two different species into different wells would not capture doublets because of the *de facto* unique indexes for each species imparted by the different wells for the first level of indexing. We therefore assumed that the assumptions that have been made and tested for standard sci-ATAC-seq and related combinatorial technologies also apply to sciMAP-ATAC, as the novel components of the workflow are in the processing prior to the combinatorial indexing stages.

With our set of 384 unique transposase indexes and the sorting of 88 nuclei per well across experiments, this would result in a doublet rate (i.e. two nuclei of the same transposase index ending up in the same PCR well) of 10.5% if the yield of sorted nuclei was perfect. However, we favor speed over precise quantification during the sorting step, as the actual number of sorted cells does not matter as long as it ends up being below the target number. We have found that using our fast sorting workflow, of the target number of events that are sorted, only between 25 and 50% are true nuclei. The rest of the events are empty droplets. We also note that these droplets do not contain ambient chromatin based on human-mouse mixing experiments^19^. Using the high end of the approximate 50% true nuclei yield, the expected doublet rate is 5.4%, in line with other commercially available single-cell platforms. When factoring in the actual yield with respect to single-cell profiles produced, the doublet rate is even lower. For example, the punch dissociation development sciMAP-ATAC preparation produced 8,012 single-cell profiles over 384 unique indexed transposition wells, for an average of just under 21 cells produced per well out of the 88 events that were sorted – a 23.7% yield. The final expected doublet rate is therefore most accurately calculated according to 21 indexed nuclei produced per well with a transposase index space of 384 for a doublet rate of 2.5%, which is well within the accepted range.

### Sequence data processing

Data analysis and plotting were performed primarily using the ‘scitools’ software (github.com/adeylab/scitools)^22^, which includes wrappers for numerous external tools. Raw sequence reads had their index combinations matched to a whitelist of expected indexed using ‘scitools fastq-dump’ which allows for a hamming distance of two and produces error-corrected fastq files. These were then aligned to mouse or human reference genome (mm10 or hg38) via bwa mem (v0.7.15-r1140)^65^ and sorted using ‘scitools align’. PCR duplicate removal and filtering for quality 10 aligned autosomal and chromosome X reads (*i.e.* excluding mitochondrial, chromosome Y, and unanchored contigs) was performed using ‘scitools rmdup’ using default parameters and plotted using ‘scitools plot-complexity’. Projections of passing reads given increased sequencing depth was performed using ‘scitools bam-project’ on the pre-duplicate removed bam file, which generates a model for every single cell based on sampling reads and calculating the passing read percentage that empirically falls within 2% accuracy^19^. Bam files were then filtered to only contain cell barcodes that contained a minimum of 1,000 passing reads and a percent unique reads less than 80 (any overly complex cell libraries may be doublets and were therefore excluded). For the human VISp dataset, cells were also filtered to have a TSS enrichment (per cell calculation) of 2 (see section “Quality metric calculations” below).

### Chromatin accessibility analysis

The filtered bam file was used for chromatin accessibility peak calling for each of the five experiments individually as well as on a combined bam file from the mouse whole brain sciATAC-seq, mouse punch dissociation development sciMAP-ATAC, and mouse SSp cortex sciMAP-ATAC experiments for the combined dataset analysis. Peak calling was run using the wrapper function ‘scitools callpeak’, which utilized macs2 (v2.1.1.20160309) for peak calling and then filtering and peak extension to 500bp^66^. Called peaks from mouse whole brain sciATAC-seq, mouse punch dissociation development sciMAP-ATAC, and mouse SSp cortex sciMAP-ATAC datasets were merged to generate a union peak set that was used to compare sciATAC-seq and sciMAP-ATAC clustering. Peak bed files and filtered bam files were then used to construct counts matrix of cells × peaks. Latent Dirichlet Allocation using the package cisTopic^67^ was performed using the scitools wrapper function ‘scitools cistopic’. Topic enrichments for region type annotations (Extended Data Figure 4g) were annotated using cisTopic function *annotateRegions*, using the Bioconductor package *TxDb.Hsapiens.UCSC.mm10.knownGene* and annotation database *org.Mm.eg.db*. The topic by annotation heatmap was plotted using cisTopic function *signaturesHeatmap*. The cells × topics matrix was biclustered and plotted using ‘scitools matrix-bicluster’, which utilizes the *Heatmap* function in the *ComplexHeatmap* package in R (v1.20.0)^68^. Two-dimensional visualization was performed using UMAP via ‘scitools umap’ and plotted using ‘scitools plot-dims’. Visualization of topic weights on the UMAP coordinates was performed using ‘scitools plot-dims’ with −M as the cells × topics matrix. Clustering was performed on the cells × topics matrix using the R package *Rphenograph*, which employs Louvain clustering and was executed using the wrapper function ‘scitools matrix-pg’ (v0.99.1)^69^. In addition to topic analyses, we utilized ChromVAR^50^ to assess the global motif accessibility profiles of cells using the wrapper function ‘scitools chromvar’ on the bam file with added read group tags using ‘scitools addrg’. Boxplots illustrating TF motif enrichment per cell were generated using values from the *Chromvar* deviations_scores matrix and plotted using *geom_boxplot* from the package *ggplot* in R, where lower and upper hinges indicate first and third quartiles, center line indicates median, upper and lower whiskers indicate 1.5 times the inner quartile range (IQR). Data points beyond the end of the whiskers are called "outlying" points and are plotted individually. All boxplot comparison significance calculations were performed used the *wilcox.test* function, indicating *paired=FALSE* and *p.adjust.method* set to Bonferroni-Holm correction in R (v0.3.0).

### Quality metric calculations

To generate tagmentation site density plots centered around transcription start sites (TSSs), we first subset filtered experiment bam files into respective annotations. We used the alignment position (chromosome and start site) for each read to generate a bed file that was then fed into the BEDOPS closest-feature command mapped the distance between all read start sites and transcription start sites (v 2.4.36, ref^70^). From this, we collapsed distances into a counts table respective of experiment and annotation and generated percentage of read start site distances within each counts table. We plotted these data using R (v 3.6.1) and *ggplot2* (v 3.3.2) *geom_line* function (default parameters) subset to 2000 base pairs around the start site to visualize enrichment. TSS enrichment values were calculated for each experimental condition using the method established by the ENCODE project (https://www.encodeproject.org/data-standards/terms/enrichment), whereby the aggregate distribution of reads ±1,000 bp centered on the set of TSSs is then used to generate 100 bp windows at the flanks of the distribution as the background and then through the distribution, where the maximum window centered on the TSS is used to calculate the fold enrichment over the outer flanking windows. The fraction of reads in a defined read set (FRiS) was used as an alternative to the fraction of reads in peaks for two major reasons. The first is that FRiP is highly dependent on the number of peaks that are called, which is, in turn, highly dependent on a) the number of cells profiled, and b) the depth of sequencing. One can increase FRiP values by sequencing a library more deeply or profiling larger numbers of cells at the same depth without reflecting any difference in underlying data quality. Second, peak calling on a population of cells favors peaks in high abundance cell types, as they make up more of the data going into the peak calling. Therefore, cells of a cell type that is lower abundance will have fewer peaks called that are specifically associated with that cell type owing to the dominance of signal by the more abundant cell type and consequently reducing the FRiP of those cells. Using FRiS instead largely avoids the challenges associated with peak calling by leveraging a comprehensive reference dataset. For the mouse FRiS calculations, we aggregated peaks that are available from mouse bulk ATAC-seq and DNAse hypersensitivity experiments provided by the ENCODE project, followed by peak collapsing, resulting in 2,377,227 total peaks averaging 744.9 bp. For the human dataset, we used a human reference dataset for DNAse hypersensitivity^28^ that contains 3,591,898 loci defined as TF footprints with an average size of 203.9 bp leading to the lower FRiS values when compared to the aggregate mouse ATAC-seq peak dataset.

### Cell type identification

The identified clusters were assigned to their respective cell type by examining the chromatin accessibility profile of marker genes that correspond to known cell types. Gene regions were plotted using ‘scitools plot-reads’ using the filtered bam file and genome track plots were generated using *CoveragePlots* from the analysis suite of tools, *Signac* (v0.2.5, https://github.com/timoast/signac). Additional support for identified cell types was performed by assessing the chromVAR results for global motif accessibility. Marker genes used for cell type identification included: *Gfap*, *Glul,* and *Agt* for astrocytes, *Col19a1* for all neuronal cell types, *Gad1, Gad2, Pvalb, Dlx1,* and *Dlx2* for GABAergic neurons, Slc17a7, *Drd1, Drd2, Bcl11b (Ctip2)* and *Ppp1r1b* for GABAergic medium spiny neurons (MSNs), also referred to as spiny projection neurons (SPNs), *C1qa, C1qc, Cx3cr1* for microglia, *Mrc1* for macrophages within the microglia cluster, *Kdr* and *Flt1* for endothelia, *Olig1* for al oligodendrocyte cell types, *Top2a* and *Cspg4 (NG2)* for OPCs, *Fyn* and *Prox1* for newly formed oligodendrocytes, and *Mobp, Mog, Cldn11* and *Prox1* for mature myelinating oligodendrocytes.

### Gene ontology enrichment analysis

Gene ontology enrichment analysis was performed for the genomic regions defined within Topic 30, the topic enriched in ischemia specific cells. Single nearest genes to Topic 30 regions were identified using GREAT (v4.0.4) for reference genome mm10^71^. Gene ontology term statistical overrepresentation for GO biological processes was calculated using Panther (v14) binomial test with false discovery rate (FDR) correction for overrepresentation of Topic 30 genomic regions in comparison to all mouse (mm10) genes. Data were plotted using *ggplot* plotting function *geom_barplot* in R (v3.2.1) with height corresponding to log_2_ GO term fold enrichment and colored by GO term −log_10_ FDR Q-value.

### Transcription factor and site enrichment through trajectories

ChromVAR analysis (described previously) was performed on all cells derived from ischemia mouse models, including the ischemic (stroke) hemisphere and contralateral (contra) hemisphere. For the cells x TF motif enrichment (TME) matrix, cells were annotated by the punch they were derived from, and a linear regression of TME as a function of punch location for each cell using the base function *lm* in R (v3.6.1). Slopes of the linear model for the ischemic and contralateral hemispheres were defined as the coefficient of the fit. The statistical significance of the interaction between TME over space and disease condition (stroke versus contralateral hemisphere) was calculated by performing an analysis of variance (ANOVA, *anova* base R v3.6.1) on the interaction of hemisphere on the linear regression defined by TF motif enrichment as a function of punch position (TME~Punch*Hemisphere_(Stroke/Contra)_), and slopes were compared using *lsmeans::lstrends* (v2.30-0). Slopes were compared between the stroke and contralateral hemispheres by taking the difference between the slopes (Δ Slope=slope_stroke_-slope_contra_). The change in slope was z-scored to center and scale TME difference, where z-score Δ Slope is equal to two standard deviations from the mean. Volcano plot of −Log_10_ P-value by Δ Slope was generated using the package EnhancedVolcano (v1.4.0) in R^72^. Line plots vignettes were generated by plotting volcano plot data using *ggplot* plotting function *geom_smooth,* method *lm* (v3.2.1). Heatmaps illustrating cell-type specific TME over space were generated by subsetting ischemia mouse model cells by cell type, and plotting TME by punch, compared between stroke and contralateral hemispheres using package ComplexHeatmap (v2.0.0) in R.

Analysis of putative regulatory elements was performed by assessing the ATAC peak probabilistic weight per cell (*cisTopic* predicitive distribution) across cells derived from punches taken from the infarct core to infarct border axis (punch positions 5 to 8) in the stroke and contralateral hemispheres, aggregated across all MCAO mice. This was performed similarly to TF motif enrichment described above, where ATAC peak probability per cell was averaged by punch position (punch positions 5-8). ATAC peak probability along the 5-8 axis was fit to a linear model and the slope in the stroke hemisphere was compared to the slope in the contralateral hemisphere in order to generate significance and delta-slope values. We found that 3,852 peaks out of 104,773 total peaks (4.8%) vary significantly across the 5-8 axis in MCAO stroke hemispheres in contrast to the contralateral hemispheres. In order to identify putative regulatory elements which are associated with the progressive gradient of glial reactivity from the infarct core to the infarct border in stroke hemipsheres, we subset our spatially significant peak set to those which uniformly increase or decrease along the 5-8 axis in stroke hemispheres. We found 72 sites which uniformly increase with increasing proximity to the infarct border, and no sites which uniformly decrease. We report all 3,852 spatially significant peaks as a reference for future MCAO regulatory element studies and denote the 72 uniformly increasing sites.

